# Beneficial Effects of Moderate Hepatic Activin A Expression on Metabolic pathways, Inflammation, and Atherosclerosis

**DOI:** 10.1101/2022.07.05.498830

**Authors:** Huan Liu, Margaret Hallauer Hastings, Robert Kitchen, Chunyang Xiao, Justin Ralph Baldovino Guerra, Alexandra Kuznetsov, Anthony Rosenzweig

## Abstract

**BACKGROUND:** Atherosclerosis is an inflammatory vascular disease marked by hyperlipidemia and hematopoietic stem cell (HSC) expansion. Activin A, a member of the Activin/GDF/TGFβ/BMP family is broadly expressed and increases in human atherosclerosis, but its functional effects *in vivo* in this context remain unclear.

**METHODS:** We studied LDLR^-/-^ mice on a Western diet for 12 weeks and used adeno-associated viral vectors with a liver-specific thyroxine binding globulin (TBG) promoter to express Activin A or GFP (control). Atherosclerotic lesions were analyzed by oil red staining. Blood lipid profiling was performed by HPLC (High Performance Liquid Chromatography), and immune cells were evaluated by flow cytometry. Liver RNA-sequencing was performed to explore the underlying mechanisms.

**RESULTS:** Activin A expression decreased in both livers and aortae from LDLR^-/-^ mice fed a Western diet compared with chow. AAV-TBG-Activin A increased Activin A hepatic expression (∼10-fold at 12-weeks, p<0.0001) and circulating Activin A levels (∼2000pg/ml vs ∼50pg/ml, p<0.001, compared with controls). Hepatic Activin A expression decreased plasma total and low-density lipoprotein (LDL) cholesterol (∼60% and ∼40%, respectively), reduced inflammatory cells in aortae and proliferating hematopoietic stem cells (HSC) in bone marrow, and reduced atherosclerotic lesion area in the aortic arch by ∼60%. Activin A also attenuated liver steatosis and expression of the lipogenesis genes, Srebp1 and Srebp2. RNA sequencing revealed Activin A not only blocked expression of genes involved in hepatic *de novo* lipogenesis but also fatty acid uptake, and liver inflammation. In addition, Activin A expressed in the liver also reduced white fat tissue accumulation, decreased adipocyte size, and improved glucose tolerance.

**CONCLUSIONS:** Our studies reveal hepatic Activin A expression reduces inflammation, HSC expansion, liver steatosis, circulating cholesterol, and fat accumulation, which likely all contribute to the observed protection against atherosclerosis. The reduced Activin A observed in LDLR^-/-^ mice on a Western diet appears maladaptive and deleterious for atherogenesis.

## Introduction

Cardiovascular diseases (CVD) are the leading cause of death worldwide^1^. Atherosclerosis is a progressive CVD characterized by accumulation of lipid plaques in arterial walls with inflammation and hematopoietic stem cell (HSC) expansion^2,3^, and often driven by increased low density lipoprotein (LDL) cholesterol^4^. Understanding better the mechanisms of atherosclerosis and identifying potential new therapeutic approaches to this condition has important clinical implications.

The liver is central to lipid metabolism including involves in uptake, esterification, oxidation, and secretion of fatty acids (FAs) with important downstream effects on systemic inflammation and metabolism. The liver is also responsible for synthesis, uptake, storage, and efflux of cholesterol, essential for multiple biological processes^5-7^. Cholesterol, triglycerides, and other lipids are assembled in the liver into lipoproteins, which are transported to peripheral tissues through the circulation^5, 6^. Hepatic free cholesterol can be esterified, incorporated into very low-density lipoprotein (VLDL) particles, secreted into the bloodstream, further converted into smaller LDL particles, and then oxidized and taken up by macrophages to form foam cells, which promote the formation of atherosclerotic plaques^5, 6^. Thus, hepatic lipid metabolism and cholesterol synthesis are essential for health of the organism but also have potential deleterious effects, such as atherosclerosis.

Hepatic sterol regulatory element binding proteins (SREBPs) play important roles in atherosclerosis. These include the transcription factors, Srebp1a, Srebp1c (alternative splicing products of Srebp1, also known as Srebf1), which are involved in synthesis of cholesterol, fatty acids, and triglycerides (TGs), and Srebp2 (Srebf2), which regulates expression of HMG-CoA (3-hydroxy-3-methylglutaryl-CoA) reductase, the target of statins, and the LDL receptor (LDLR)^8^.

Hepatic Srebp1c expression increases atherogenic lipoprotein profiles and accelerates atherosclerosis in LDLR^-/-^ mice^9^. Concordantly, an inhibitor of Srebp processing reduces atherosclerosis^10^. Similarly, GPR146 deficiency reduces hepatic Srebp2 activity and protects against atherosclerosis^11^.

Activin A is a member of Activin/GDF/TGFβ/BMP protein family. Many members of this family, including Activin A, drive a metabolic switch from an anabolic to catabolic state. Activin A signals through type I (ACTR1B [ALK4], ACTR1C [ALK7] or TGFBR1) and type II (ACVR2A, ACVR2B, and BMPR2) receptors to induce Smad2/3 phosphorylation and nuclear translocation regulating gene expression^12, 13^. Follistatin (FST) is induced by Activin signaling and antagonizes its ACVR receptor binding site^14^, forming a negative feed-back loop^15^. Activin A can also form a non-signaling complex with type I (ACVR1A, ALK2) and type II receptors^16-18^.

The role of Activin A in atherosclerosis is unclear. On one hand, Activin A is expressed in the intima in early atherosclerotic lesions and elevated further in lesions of advanced atherosclerotic patients^19, 20^. Plasma Activin A is increased in angina patients (∼500 pg/ml) compared with controls (∼300 pg/ml)^19-21^. These observations suggest Activin A could contribute to atherosclerosis progression. On the other hand, *in vitro* studies demonstrate that Activin A inhibits foam cell formation in Thp1 and Raw 264.7 macrophages^22, 23^, thought to model an early step in atherogenesis, while FST promotes foam cell formation in Thp1 macrophages^24, 25^. Exogenous Activin A has also been reported to reduce neointimal hyperplasia in a femoral artery cuff murine model *in vivo*^26^. These studies suggest Activin A may be anti-atherogenic and focus on its role in vessel wall constituents. Thus, we lack an understanding of the *in vivo* effects of Activin in atherosclerosis and the role of different sites of expression.

Here we find that Activin A is highly expressed in aorta and liver but decreases in both in LDLR^-/-^ mice fed a Western diet compared with those fed chow. Hepatic Activin A expression mediated by an adeno-associated viral vector in LDLR^-/-^ atherosclerotic mice increased circulating Activin A levels (∼2000 pg/ml vs ∼50pg/ml, p<0.0001) and protected against atherosclerosis by reducing cholesterol, decreasing inflammation, and reducing proliferating HSCs. Mechanistically, hepatic Activin A reduced liver Srebp1/Srebp2 expression, pathways involved in hepatic lipogenesis and fatty acid uptake, as well as liver steatosis. In addition, hepatic Activin A reduced body fat accumulation and improved glucose tolerance. We conclude that the reduction in hepatic Activin A observed on a Western diet is maladaptive and could represent an evolutionary vestige perhaps valuable in calorie-poor environment but potentially maladaptive amidst caloric abundance. Whether AAV8-TBG-Activin A has therapeutic potential in atherosclerosis and related metabolic diseases may warrant further investigation.

## Methods

The data, methods, and study materials used to conduct the research will be available from the corresponding author upon reasonable request.

### Animal Studies

All mouse experiments were performed in accordance with the NIH Guide for the Care and Use of Laboratory Animals and approved by the Institutional Animal Care and Use Committee (IACUC) of Massachusetts General Hospital.

For the atherosclerosis model, 8-week-old male LDLR^-/-^ (#002207, body weight ∼24-25g) obtained from Jackson Laboratory were injected via tail vein with AAV8-TBG-GFP or AAV8-TBG-Activin A viruses at 2×10^10^ genome copies/mouse. After 1 week, mice were fed a high cholesterol Western diet (Envigo Teklad, TD.96121) or standard chow diet for 12 weeks. Mice were fasted overnight before euthanasia.

A detailed description of the methods and supporting data are available in the Online Supplemental materials.

### Statistical Analysis

Data are expressed as mean±SEM and analyzed using GraphPad Prism 8 (GraphPad Software) by unpaired Student t-tests. p<0.05 was considered significant. *p<0.05, **p<0.005, and ***p<0.001.

## Results

### Activin A mRNA decreases in aorta and liver in LDLR^-/-^ mice fed a Western diet

We examined plasma Activin A levels as well as the expression of the Activin A transcript in liver and aorta, and how these were affected by hyperlipidemia. Plasma Activin A levels were increased in LDLR^-/-^ mice fed a Western diet compared with chow (Figure 1A) while plasma Follistatin (FST) levels were similarly undetectable in all mice (data not shown). Baseline Activin A mRNA levels measured by QPCR were high in aorta and liver relative to other organs tested (Figure S1A). Surprisingly, given previous reports of increased Activin A expression in atherosclerosis^19-21^, Activin A mRNA levels decreased in both aortae and livers from LDLR^-/-^ mice fed a Western diet compared with chow (Figure 1B-1C). In contrast, FST was expressed most highly in muscle and aorta, and at lower levels in liver (Figure S1B) and was unaffected by diet (Figure 1B-1C). Consistent with these observations, we found that oxidized LDL (oxLDL) decreased Activin A but not FST mRNA in primary peritoneal macrophages and the Hepa 1-6 hepatic carcinoma cell line (Figure 1D-1E). Together these observations suggest the Activin pathway is suppressed in the liver and aorta, and this may be a direct effect of oxLDL.

**Figure 1.**
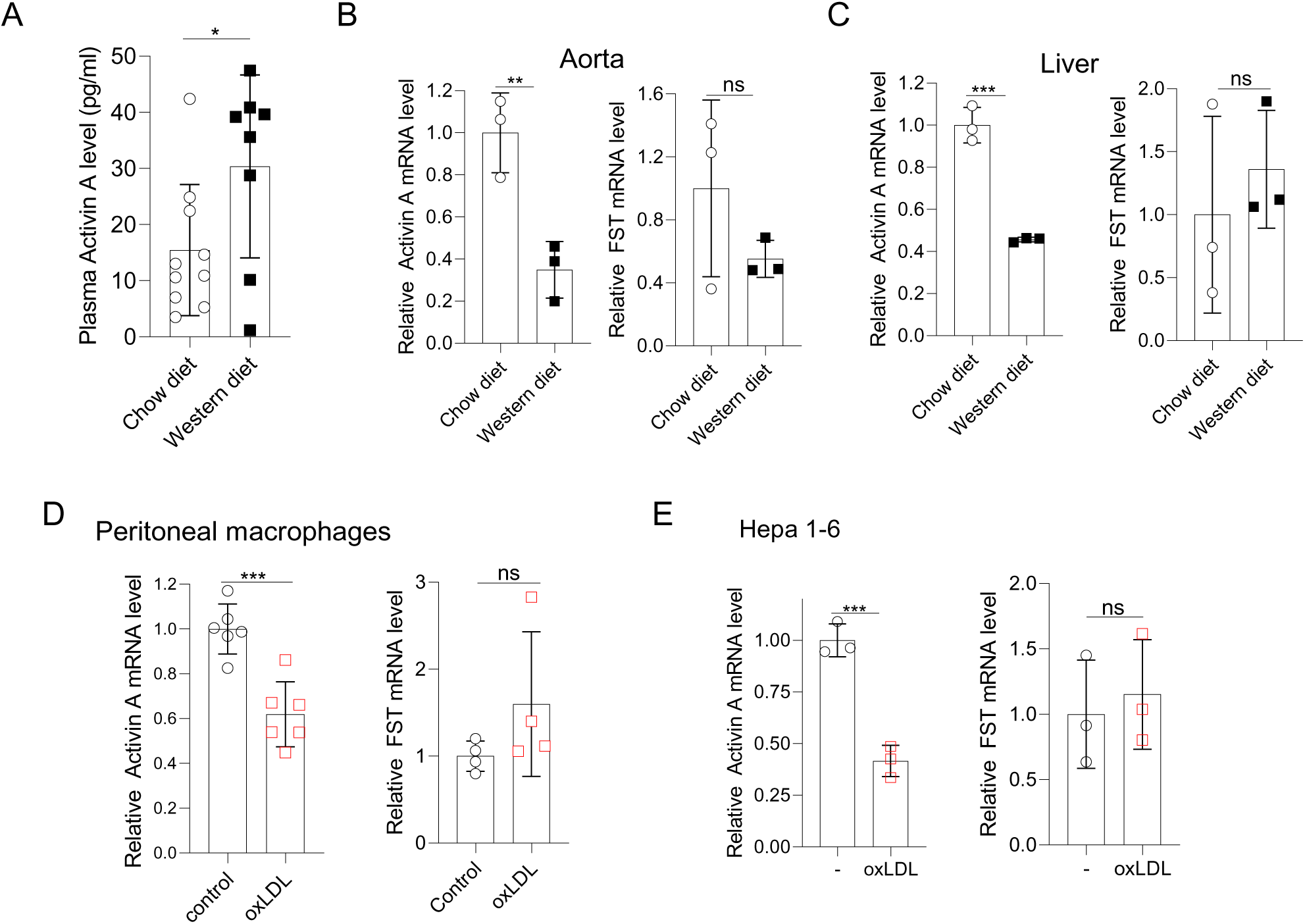
Activin A mRNA decreases in aorta and liver in western diet fed mice, oxLDL treated macrophages and Hepa 1-6 cells. **A**. ELISA shows plasma Activin A level in LDLR^-/-^ mice fed with western diet for 12 weeks compared to chow diet. Plasma FST levels are close to baseline (0) and not shown here. N=8-10 mice. **B-C**. Q-PCR shows the Activin A but not FST mRNA decreases in the aorta and liver in LDLR^-/-^ mice fed the western diet for 12 weeks compared to chow diet. N=3 mice. **D**.Q-PCR shows Activin A but not FST mRNA decreases in peritoneal macrophages after oxLDL treatment. Peritoneal macrophages were treated with and without 10μg/ml oxLDL in 0.5%BSA (fatty acid free)/RPMI at 37°C for 24h. **E.** Q-PCR shows Activin A but not FST mRNA decreased in oxLDL treated Hepa 1-6 cells. Hepa 1-6 cells were treated with and without 10 μg/ml oxLDL in 0.5%BSA (fatty acid free)/RPMI 1640 at 37°C for 24h.

### Activin A protects against atherosclerosis *in vivo*

We generated AAV8 vectors using the TBG promoter to drive liver-specific^27, 32-34^ expression of Activin or GFP. We confirmed TBG-driven Activin A mRNA expression in HepG2 cells and protein in the media, as well as induction of Smad2/3 phosphorylation that was blocked by FST-315, the circulating isoform of FST (Figure S2A). Similar to recombinant Activin A protein^24, 25^, Activin A from HepG2 conditioned media also reduced oxLDL uptake by peritoneal macrophages *in vitro* (Figure 2A-2B). Thus, the AAV8-TBG-Activin A vector effectively induces Activin A expression and functional protein secretion.

**Figure 2.**
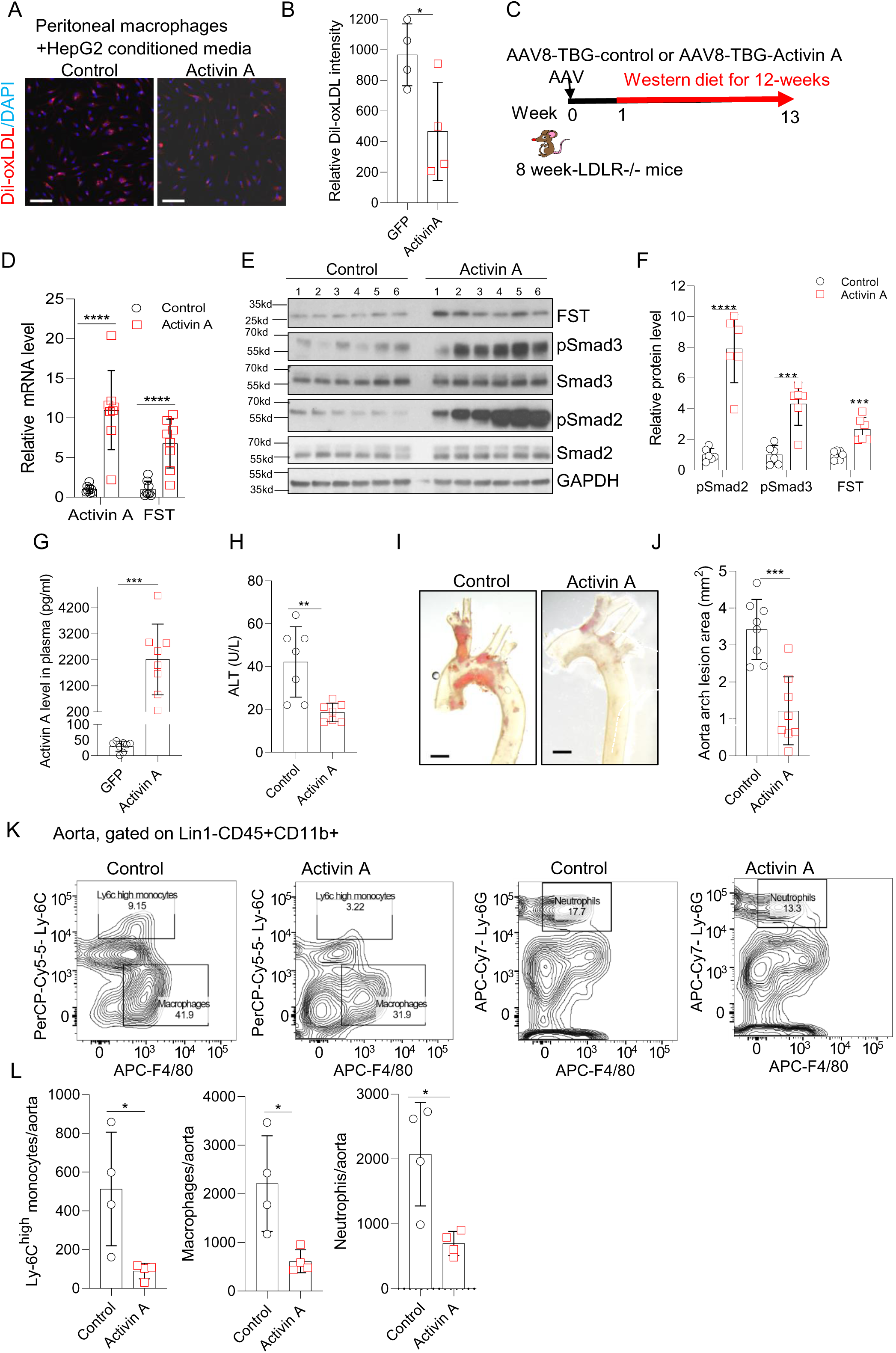
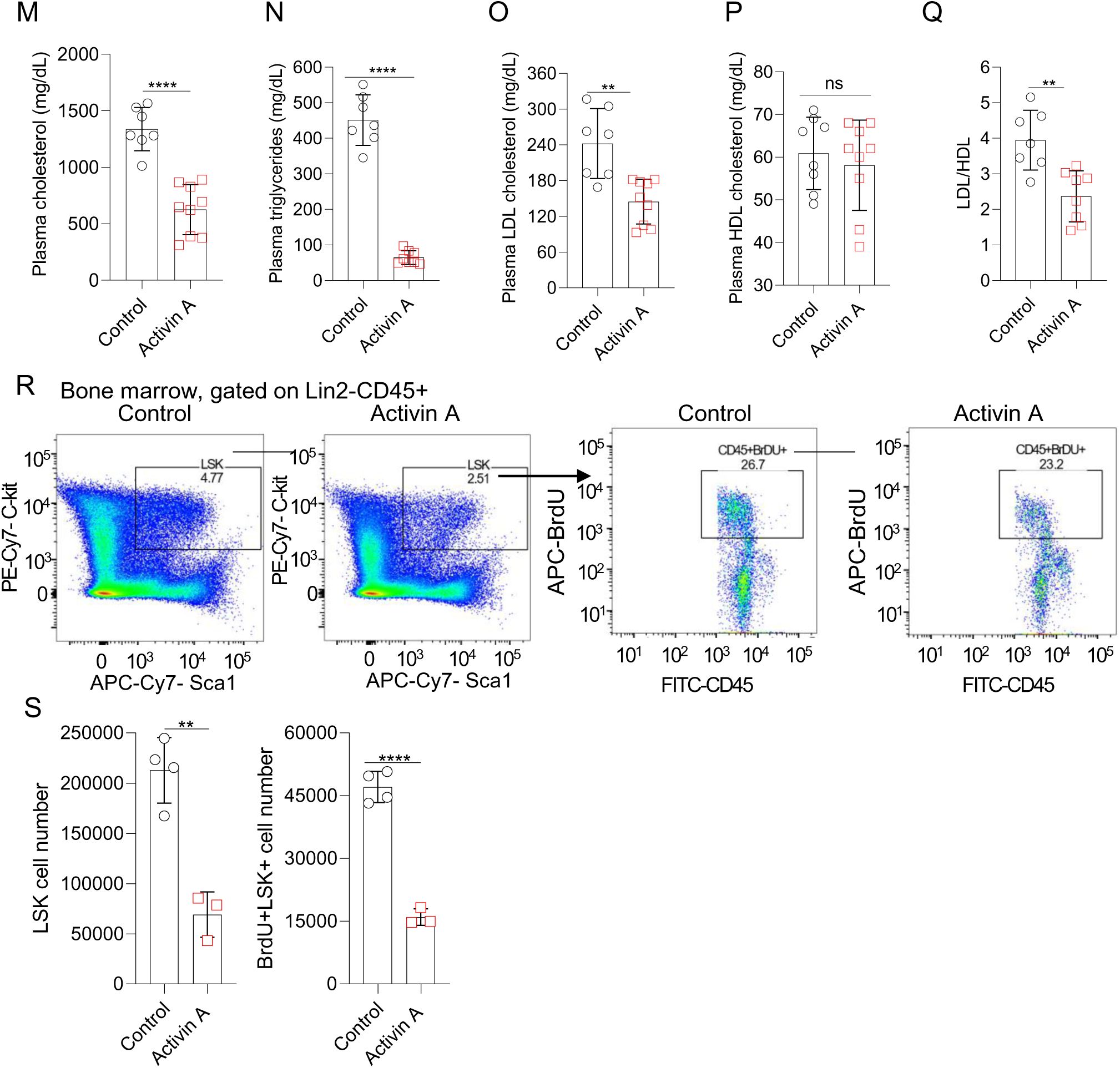
Activin A protects against atherosclerosis *in vivo* and reduces proliferating hematopoietic stem cells number. **A**. Confocal images of the activity of Peritoneal macrophages oxLDL uptake decreases after Activin A expressed Hepg2 conditioned media treatment compared to control. **B**. Graph represents the quantification of the mean Dil-oxLDL intensity/cell by image J. **C**. Schematic of the experimental design for the atherosclerosis model. 8-week-old male LDLR^-/-^ mice are delivered 2×10^10 genome copies of AAV8-TBG-control (control) or AAV8-TBG-Activin A (Activin A) viruses by tail vein injection, then after 1 week fed the Western diet for 12 weeks to induce Atherosclerosis **D**. Q-PCR shows liver Activin A, and its direct target gene, FST mRNA were induced in Activin A treated atherosclerotic LDLR^-/-^ mice compared with control. N=8 mice. **E**. Western blotting shows the Activin signaling pathway is activated in Activin A expressing liver compared with control. Antibodies for Smad2, pSmad2, Smad3, pSmad3 and the Activin signaling pathway direct target gene FST were used, and GAPDH as the loading control. **F**. Quantification of relative pSmad2, pSmad3, and FST protein level. **G**. ELISA shows Activin A protein circulating in the plasma of atherosclerotic LDLR^-/-^ mice after AAV8-TBG-Activin A treated and western diet for 12 weeks. N=8 mice. **H.** Graph represents plasma ALT (SGPT) decreases in Activin A treated atherosclerotic mice compared with control. **I.** Images show the oil red staining of atherosclerotic plaques in the aortic arch in control and Activin A expressing atherosclerotic mice. Scale bar, 1mm. **J**. Quantification of lesion area in the aortic arch. N=8 mice. **K.** Assessment and gating strategy of aortic immune cells by flow cytometry. Lin1: Ter119/NK1.1/CD19/CD90.2/CD3. Ly-6C^high^monocytes (CD45+Lin1−CD11b+F4/80−Ly-6C^high^), macrophages (CD45+Lin1−CD11b+F4/80+Ly-6C^low^) and neutrophils (CD45+Lin1−CD11b+Ly-6G+F4/80−). **L**. Quantification of Ly-6C^high^ monocytes, macrophages, and neutrophils in aorta. N=4 mice. **M-Q.** Graphs represent the plasma lipid profiles including cholesterol, triglycerides, LDL, HDL and LDL/HDL in control and Activin A-expressing atherosclerotic mice. N=7-9 mice. **R.** Assessment and gating strategy of hematopoietic stem cells (HSC) proliferation by flow cytometry. Lin2: CD3/Ly-6G/Ly-6C/ B220/CD11B/Ter119/CD127. LSK: Lin2- CD45+Sca1+c-Kit+. **S.** Quantification of the of LSK cells and BrdU+ LSK cells. HSC were from 2 tibia and 2 femurs of each atherosclerotic mouse. 1mg BrdU was injected into mice 2 hours before euthanasia. N=3-4 mice. Similar results were obtained from 3 independent experiments.

*In vivo* studies confirmed that TBG-driven GFP expression was specific to the liver (Figure S2B) and TBG also mediated hepatic Activin A expression that induced Smad2/3 phosphorylation, and increased circulating plasma Activin A (Figure S2C-S2E). LDLR^-/-^ mice were injected with AAV8-TBG-GFP (control) or AAV8-TBG-Activin A, and then fed a western diet for 12 weeks to induce early-stage atherosclerosis (Figure 2C). Hepatic Activin A expression induced Smad2/3 phosphorylation and expression of the downstream target gene FST in the liver but not aorta (Figure 2D-2F and S2F). Notably, Activin receptors, ACVR1b and ACVR2b, dramatically decreased and ACVR2a showed a nonsignificant trend toward decreasing in Activin-expressing mice compared with controls (Figure S2G). Plasma Activin A increased in AAV8-TBG-Activin A treated mice compared to controls (Figure 2G). Of note, plasma Activin A levels in control mice fed the Western diet were higher than chow diet (Figure 1A). However, plasma Activin A levels seen in Activin A-expressing mice fed a Western diet were even higher (Figure 2G). The molecular weight of the FST band detected in the liver (<35kd) was smaller than FST315 (>35kd), suggesting it may correspond to the FST288 isoform, which is limited to tissue and not secreted into the circulation. Indeed, plasma FST levels remained at background levels (data not shown), suggesting the ratio of plasma Activin A/FST was increased in the Activin A expressing mice compared with controls. Surprisingly, Activin A did not induce liver injury and plasma ALT (or SGPT for Serum Glutamic-Pyruvic Transaminase) decreased in Activin-expressing mice compared with controls (Figure 2H).

Oil red staining demonstrated that Activin A expression dramatically decreased atherosclerotic plaques in the aortic arch by ∼60% (Figure 2I-2J). In addition, flow cytometry revealed that Ly-6C^high^ monocytes, F4/80+ macrophages and Ly-6G+ neutrophils were all substantially decreased in aortae from Activin A-expressing mice compared with controls (Figure 2K-2L), suggesting both lesion size and inflammation were reduced by Activin A. Hepatic Activin A expression dramatically decreased plasma total cholesterol (>50%), LDL (∼40%), and triglycerides (>80%) (Figure 2M-2O), without altering HDL (Figure 2P); thus, reducing the LDL/HDL ratio (Figure 2Q). Taken together, hepatic Activin A protected against atherosclerosis *in vivo* and this was associated with a substantial decline in circulating atherogenic lipid levels (LDL-C and TGs) without a change in HDL or increased liver injury.

We next used flow cytometry to investigate the effect of Activin A on HSCs. Activin A decreased the absolute number of bone marrow LSK (Lin-CD45+Sca1+c-Kit+) cells, the population that includes HSCs^35^, by ∼60% (Figure S2H-S2I). In the LSK population, long-term HSCs (LtHSC, CD135-/CD150+/CD48- LSK(Lin-Sca1+c-Kit+)) differentiate into short-term HSCs (StHSC, CD135-/CD150-/CD48-LSK) and then multipotent progenitor cells (MPPS) including MPP2 (CD135-/CD150+/CD48+LSK), MPP3 (CD135-/CD150-/CD48+ LSK) and MPP4 (CD135+/CD150-/CD48+/- LSK)^36^. There was an >80% decrease in LtHSC and StHSC and ∼60% decrease in MPP2, MPP3 and MPP4 cells in atherosclerotic mice expressing Activin A compared with controls (Figure S2H-S2I).

To determine whether the decrease in LSK cells is due to an effect on HSC proliferation, we assessed BrdU incorporation by flow cytometry. LSK cells were dramatically decreased overall and BrdU+ LSK cells were reduced ∼60% in Activin A-expressing atherosclerotic mice versus controls (Figure 2R-2S). Thus, Activin A decreased the actively proliferating and the total number of HSC in atherosclerotic mice.

### Activin A decreases liver steatosis and inflammation in atherosclerotic mice

We next investigated the effects of AAV8-TBG-Activin A on the liver itself. The accumulation of excessive lipid droplets is a pathological change known as hepatic steatosis7. H&E and oil red staining revealed accumulation of much smaller lipid droplets in Activin A-expressing livers compared with controls (Figure 3A-3B). The accumulation of free cholesterol contributes to liver injury, ER stress, mitochondrial dysfunction, fibrosis inflammation, as well as cholesterol crystallization in lipid droplets and consequent hepatocyte cell death^37-40^. Activin A expression decreased hepatic total and free cholesterol by 30% without changing triglycerides, a common and “safe storage” for fatty acids (Figure 3C-3E). In addition, QPCR demonstrated that Activin A expression reduced expression of Srebp1 and Srebp2 (Figure 3F), suggesting that Activin A could inhibit hepatic lipogenesis. Picrosirius Red staining (collagen I and III) demonstrated that fibrosis was primarily located in lipid droplet-rich regions in control livers but around veins in Activin A expressing livers (Figure 3A). There was no difference in overall fibrotic area in the two groups (Figure 3H). Although the Activin A pathway was activated, as indicated by increased FST protein and Smad2/3 phosphorylation (Figure 2E-2F), fibrosis marker genes Col1a1, Col1a2, Col3a1, and MMP12 were significantly decreased (Figure 3G). In addition, there was a trend toward decreased expression of the hepatic stellate cell marker gene, Acta2, in Activin A expressing livers compared with controls (Figure 3G). Immunohistochemical staining for CD45 showed that inflammatory cells were dramatically decreased in Activin A-expressing livers compared with control (Figure 3A and 3I). Furthermore, mRNA levels of inflammatory marker genes TNFα and F4/80 were decreased in Activin A-expressing liver versus controls (Figure 3J). Collectively, these findings suggest that Activin A decreased liver steatosis and inflammation without inducing fibrosis in atherosclerotic mice.

**Figure 3.**
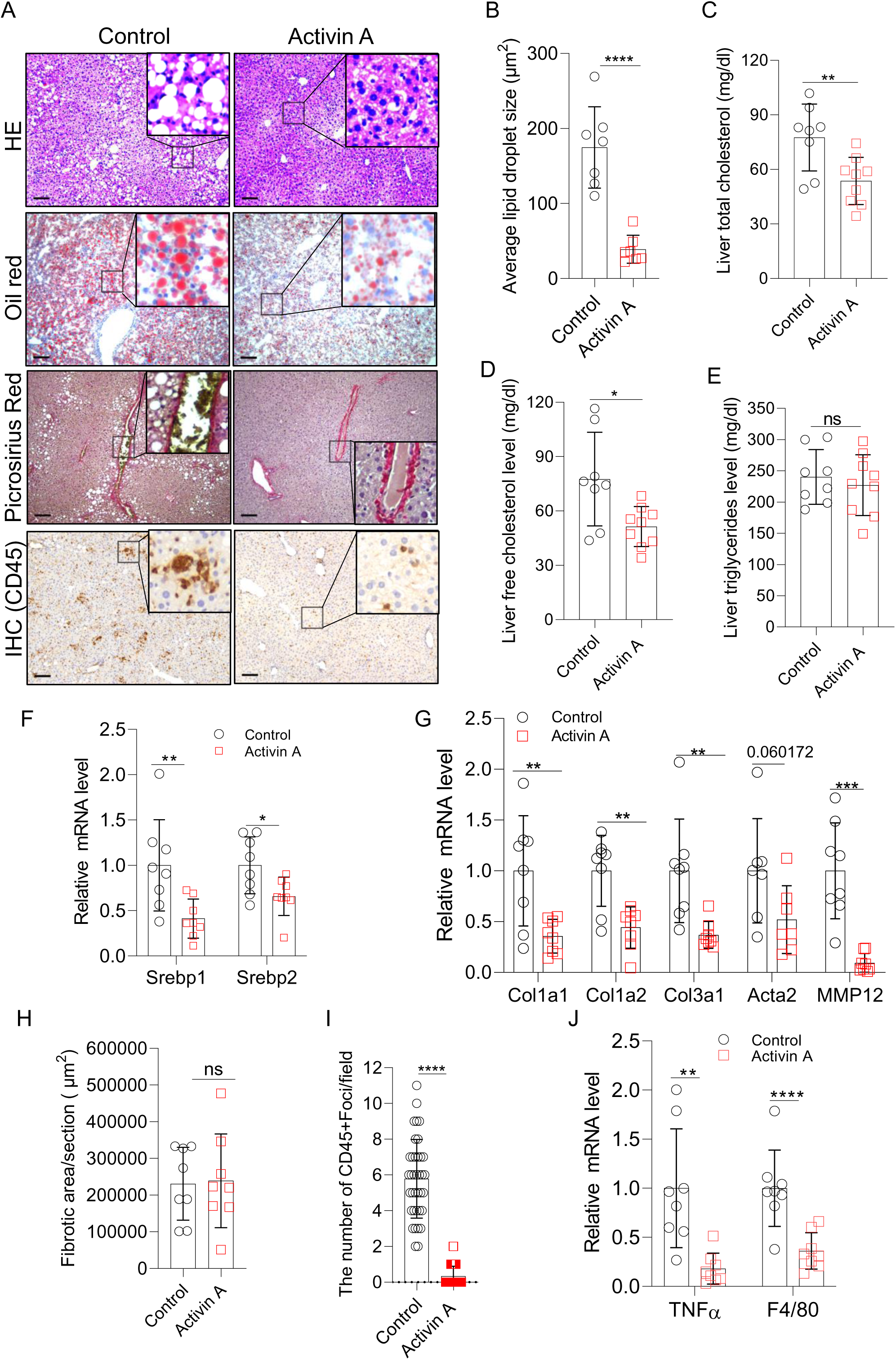
Activin A decreases liver steatosis and inflammation in atherosclerotic mice. **A**. Representative Hematoxylin and Eosin staining, oil red staining (lipid), picrosirius red staining (collagen I and III), and immunohistochemistry staining of leukocytes (with anti-CD45 antibody) in liver samples from control and Activin A treated –atherosclerotic mice. N=7-8 mice in each staining. **B**. Quantitation of the lipid droplet size based on oil red staining in A. N=8 mice. **C-E**. Graph represent the liver total cholesterol, free cholesterol level and triglycerides in atherosclerotic mice. N=7-9 mice. **F**. Q- PCR showing the mRNA level of Srebp1 and Srebp2, the key transcription factors for cholesterol synthesis and triglycerides in control and Activin A- expressing liver. N=8 mice. **G**. Q-PCR showing the mRNA level of fibrosis marker genes (Col1a1, Col1a2, Col3a1 and MMP12) and hepatic stellate cell marker gene (Acta2) in control and Activin A-expressing liver. N=8 mice. **H**. Quantification of fibrotic area of in the right lateral lobe based on picrosirius red staining in A. N=8 mice. **I**. Quantification of CD45+ foci based on IHC staining in A. CD45+ foci was defined as the sites with > 8 CD45 positive cells. The number of CD45+ foci/field (7-8 field/mice) in the right lateral lobe was quantified. N=8 mice. **J**. Q-PCR showing the mRNA level of inflammatory marker genes TNFα, and F4/80 in control and Activin A expressing- atherosclerotic liver. N=8 mice. Mean values for each mouse were used for quantification.

### Activin A reduced expression of pathways involved in fatty acid uptake, *de novo* lipogenesis and lipid associated macrophage recruitment

To identify the mechanisms responsible for the effects of hepatic Activin A, we performed RNA-sequencing in Activin A-expressing compared with control liver samples. Differential expression analysis confirmed upregulation of Activin A (also known as Inhba) and its target gene FST in Activin A-expressing mice compared with controls (Figure 4A). Overrepresentation analysis of the differentially expressed genes (DEG) identified “Fatty acid metabolic process” ranks 1 among the GO Biological Process and “lipid and atherosclerosis” ranks 8 among the KEGG pathways, and (Figure 4B and S3A), likely contributing to the decreased lipid accumulation seen in Activin A expressing livers.

**Figure 4.**
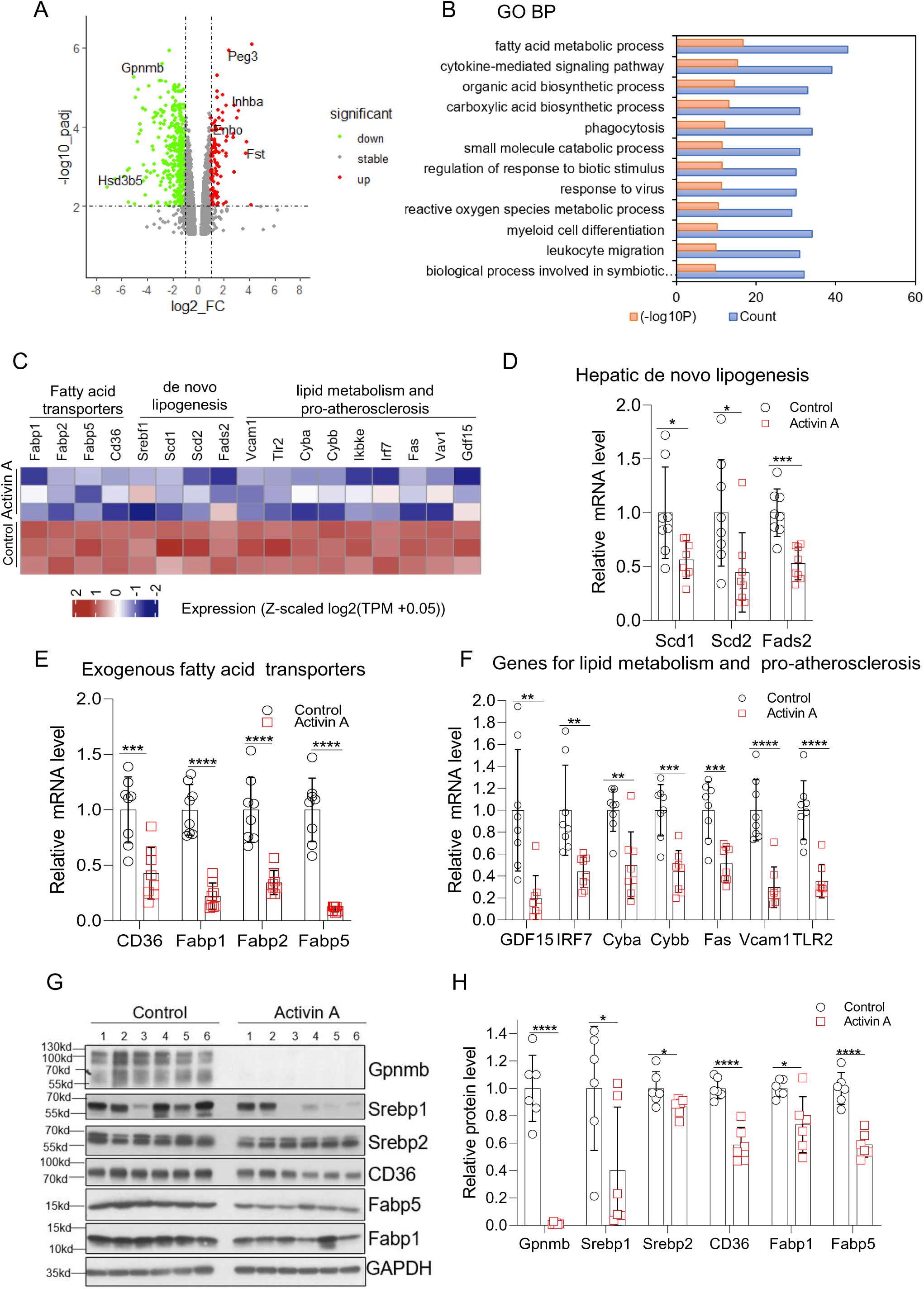
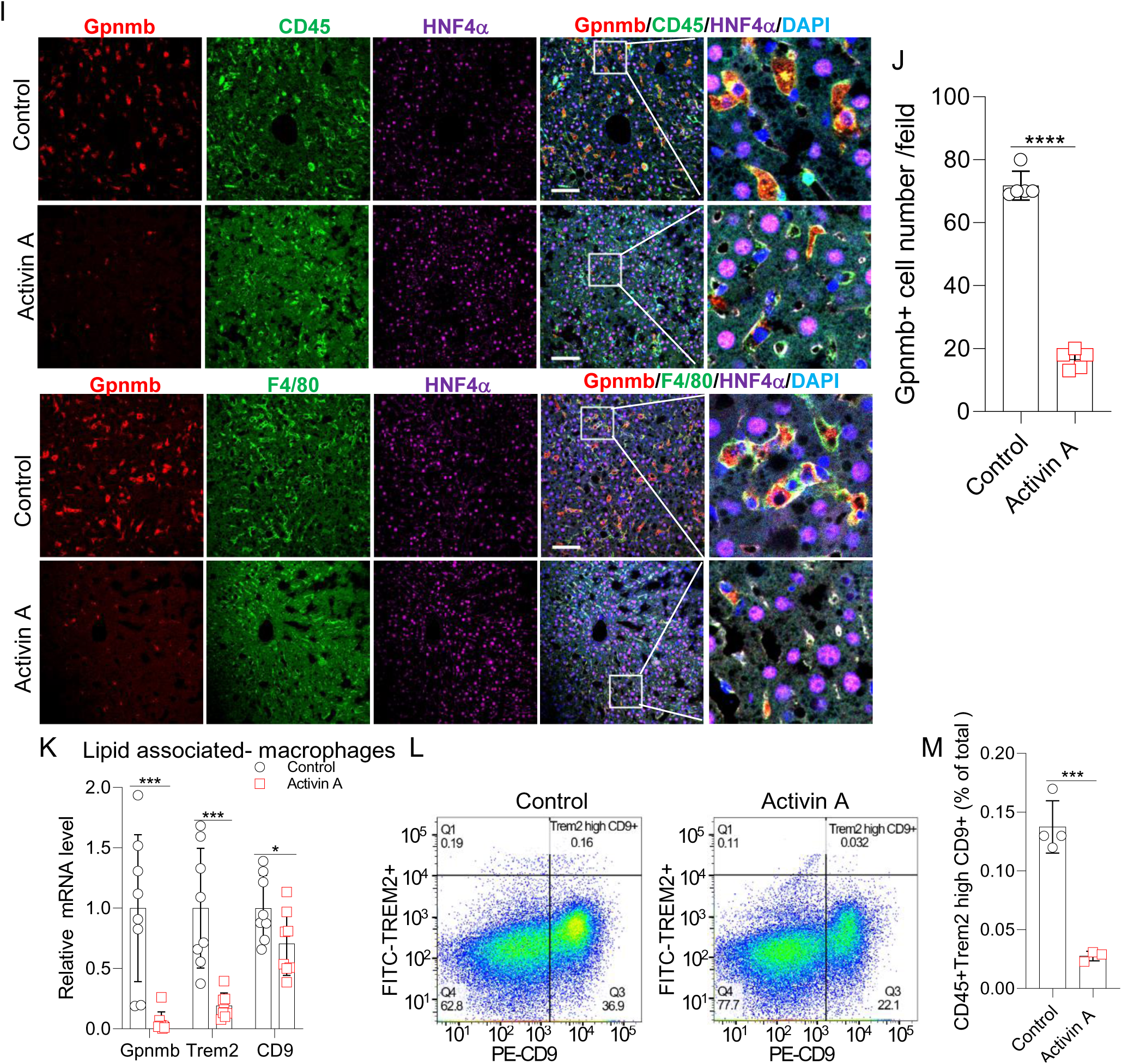
Activin A decreases expression of genes involved in fatty acid uptake, and *de novo* lipogenesis, and lipid associated- macrophage markers in atherosclerotic liver. **A.** Representative volcano plot of differentially regulated genes in RNA-seq analysis of Activin A expressed liver compared with control. Triplicates per group. TPM (control) or TPM (Activin A) ≥ 5. Significant: |log2Fc|>=1, padj<0.01. Downregulation: green, upregulation: red, and stable: grey. Labeling genes, padj<0.01, TPM>=10, log2_Fc>=3 or log2_Fc<=-5. **B**. Top 12 GO biological processes dramatically changed in RNA-seq analysis of Activin A-expressing liver compared with control, based on overrepresentation analysis. Cluster Profiles package was used. TPM (control) or TPM (Activin A) ≥ 5, p<0.05, |log2Fc|>=1. **C.** Heatmap shows differentially regulated featured genes in fatty acid metabolic process, lipid metabolism and atherosclerosis. Gene list is from GO and KEGG enrichment. TPM (control) or TPM (Activin A) ≥ 5, p<0.05, |log2Fc|>=1, padj<0.01. Relative gene expression values (Z-scaled log2(TPM +0.05)) were represented. **D-E.** Q-PCR confirmation of decreased transcript expression of genes involved in hepatic *de novo* lipogenesis and exogenous fatty acid uptake in Activin A-expressing liver compared with control. N=8 mice. **F.** Q-PCR confirmation of decreased transcript expression of genes involved in lipid metabolism and atherosclerosis in Activin A expressing liver compared with control. N=8 mice. **G**. Western blotting showing decreased Gpnmb, Srebp1, Srebp2, CD36, Fabp1, Fabp5 protein in Activin A- expressing liver compared with control. Antibodies for Gpnmb, Srebp1, Srebp2, CD36, Fabp1, and Fabp5 antibodies were used, GAPDH as the loading control. **H.** Quantification of relative Gpnmb, Srebp1, Srebp2, CD36 and Fabp5 protein level. **I.** Images show the immunofluorescence staining of Gpnmb colocalization with CD45+ and F4/80+ cells, but not hepatocytes in atherosclerotic liver. CD45 (green), leukocytes marker, F4/80 (green), macrophages marker, Gpnmb (red), hepatocyte marker, HNF4α, (purple, locates in nuclei), and nuclear stain DAPI (blue). Scale bar, 100μm. **J**. Quantification of Gpnmb+ cells in I. 4-5 random fields /section. N=5 mice. **K**. Q-PCR shows decreased mRNA levels of lipid-associated-macrophages markers Trem2, Gpnmb and CD9 in Activin A-expressing liver compared with control. N=8 mice. **L-M**. Assessment, gating strategy and quantification of lipid associated- macrophages (Trem2 high CD9+). Gpnmb antibodies did not work for flow. N=3-4 mice.

A heatmap generated based on the DEG list from the “fatty acid metabolic process” GO term and “lipid and atherosclerosis” KEGG pathway revealed decreases in multiple genes related to *de novo* lipogenesis and fatty acid uptake in Activin A-expressing livers (Figure 4C). Srebp1 is a pivotal transcription factor for triglyceride metabolism and *de novo* lipogenesis, which directly activates the key desaturases for the biosynthesis of monounsaturated fatty acids (MUFA) and polyunsaturated fatty acid (PUFAs), including Scd1(stearoyl-CoA desaturase 1, primarily in liver), Scd2 (stearoyl-CoA desaturase 2), and fatty acid desaturase (Fads2) expression^41-43^. By QPCR, we verified that Scd1/2 and Fads2 were significantly decreased in Activin A-expressing livers compared with controls (Figure 4D). FABPs are essential for intracellular binding and transport of fatty acids, as well as cholesterol and phospholipid metabolism^44^. Fabp1 is the major hepatic FABP, and its deficiency attenuates both diet-induced hepatic steatosis and fibrogenesis^45^. mRNA levels of Fabp1, Fabp2, Fabp5, and the atherogenic scavenger receptor and fatty acid transporter, CD36^46, 47^, were all decreased in Activin A-expressing liver compared with controls (Figure 4E), as were the corresponding protein levels (Figure 4G-4H). Consistently, Activin A also inhibits Srebp1, Srebp2, Fabp5 in Hepa 1-6 cells with oxLDL treatment *in vitro*. (Figure S3B). Oleic acid (OA, a monounsaturated omega-9 fatty acid) is a potent inducer of triglyceride synthesis and storage. Activin A reduced OA-induced -fatty acid uptake in Hepa 1-6 cells *in vitro*, while this effect was rescued with a neutralizing Activin A antibody (Figure S3C-S3D). Multiple differentially-regulated genes have been reported to play roles in lipid metabolism or to promote atherosclerosis, such as GDF15^48, 49^, IRF7^50^, Cyba^51^, Cybb^52^, Fas^53, 54^, Vcam1^55^ and TLR2^56^ (Figure 4C). Expression of these genes was dramatically reduced in Activin A-expressing liver (Figure 4F). Thus, Activin A likely blocks both *de novo* lipogenesis and fatty acid uptake to protect against liver steatosis, free cholesterol accumulation, and associated atherosclerosis (Figure 6F).

A recent report showed that adenoviral expression of the glycoprotein NMB (Gpnmb, also known as DC-HIL) in hepatocytes promotes lipogenesis in white adipose tissue (WAT) and exacerbates obesity and insulin resistance^57^. Gpnmb is also a core component of Trem2^high^CD9+ lipid associated- macrophages (LAM) in the liver and is induced in trans-fat containing AMLN (amylin liver non-alcoholic steatohepatitis, NASH) diet-induced nonalcoholic steatohepatitis (NASH) liver^58^, while Elafibranor, an agonist of PPARα and PPARδ, reverses NASH and dramatically decreases Gpnmb protein levels, as well as Trem2, CD9, and Gpnmb mRNA levels in mice^58^ Interestingly, hepatic Gpnmb mRNA was decreased in Activin A-expressing mice as (Figure 4A). Interestingly, we found that hepatic Gpnmb mRNA and protein expression were induced by the Western diet (Figure S3E-S3F) and dramatically decreased in Activin A-expressing mice (Figure 4A, 4G-4H, 4K). Activin A also attenuated Gpnmb expression and secretion in Hepa 1-6 cells *in vitro* (Figure S3G), suggesting this is a direct effect. Immunofluorescent staining showed that Gpnmb was colocalized with CD45+ and F4/80+ but not hepatocytes, confirming Gpnmb expression in a macrophage lineage (Figure 4I-4J). Notably, our RNA-seq data also indicated downregulation of Trem2 (Figure 5A). We next focused on LAM. The LAM marker genes Trem2 and CD9 are also decreased in Activin A-expressing livers compared with controls (Figure 4K). Moreover, we confirmed that Activin A expression led to a >3-fold decrease in LAM Trem2^high^ CD9+ populations in the liver by flow cytometry (Figure 4L-4M). Together, these data suggest that Activin A expression decreased genes involved in fatty acid uptake, *de novo* lipogenesis and lipid- associated- macrophage recruitment in atherosclerotic liver, and thus decreased liver steatosis.

**Figure 5.**
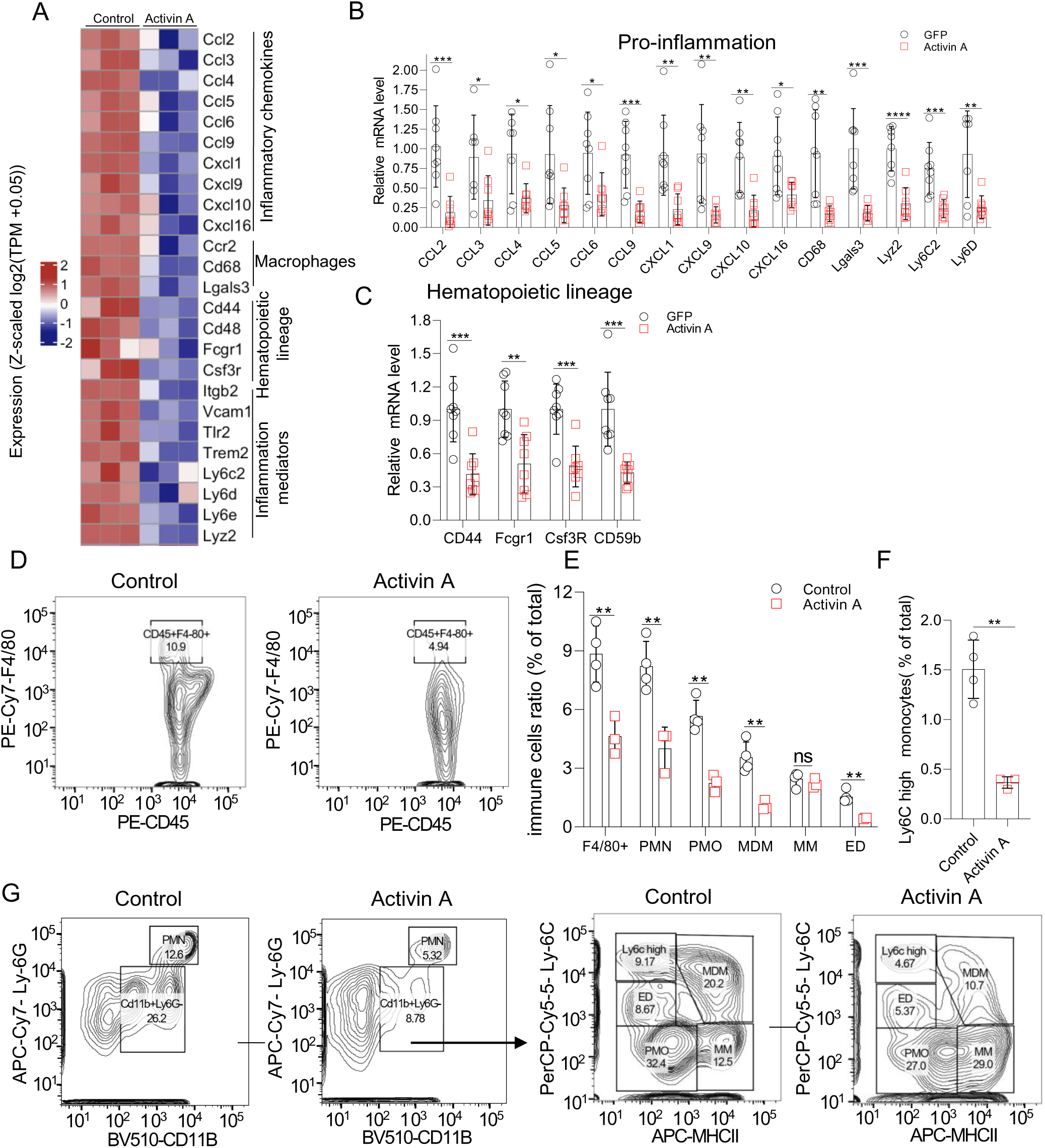
Activin A decreases liver macrophages, Ly-6C^high^ monocytes and polymorphonuclear cells (PMN) infiltration in Atherosclerotic mice. **A.** Heatmap shows differentially regulated featured genes in leukocytes migration pathway. Gene list is from GO enrichment. Relative gene expression values (Z-scaled log2(TPM +0.05)) for the 25 genes related to pro-inflammation. TPM (control) or TPM (Activin A) ≥ 5. p<0.05, |log2Fc|>=1, padj<0.01. **B-C**. Q-PCR confirmation of RNA-seq data in A. **D.** Assessment of CD45+F4-80+ populations by flow cytometry. **E-F**. Quantification of immune cells from D and G. PMN (polymorphonuclear cells, CD45+CD11b+Ly-6G+, including neutrophils, basophils, and mast cells), PMO (patrolling monocytes, CD45+CD11b+ Ly-6G-Ly-6C- MHC-II-), MDM (monocytes-derived macrophages, CD45+CD11b+ Ly-6G-Ly-6C^high^ MHC-II high), MM (mature macrophages, also known as Kupffer cells, CD45+CD11b+ Ly-6G-Ly-6C^neg/low^ / MHC-II^high^), ED (eosinophils), Ly-6C^high^ monocytes (CD45+CD11b+ Ly-6G-Ly-6C^high^/MHCII-). N=3-4 mice. Repeat for 3 times. **G**. Assessment and gating strategy of immune cells in atherosclerotic liver by flow cytometry in E and F.

**Figure 6.**
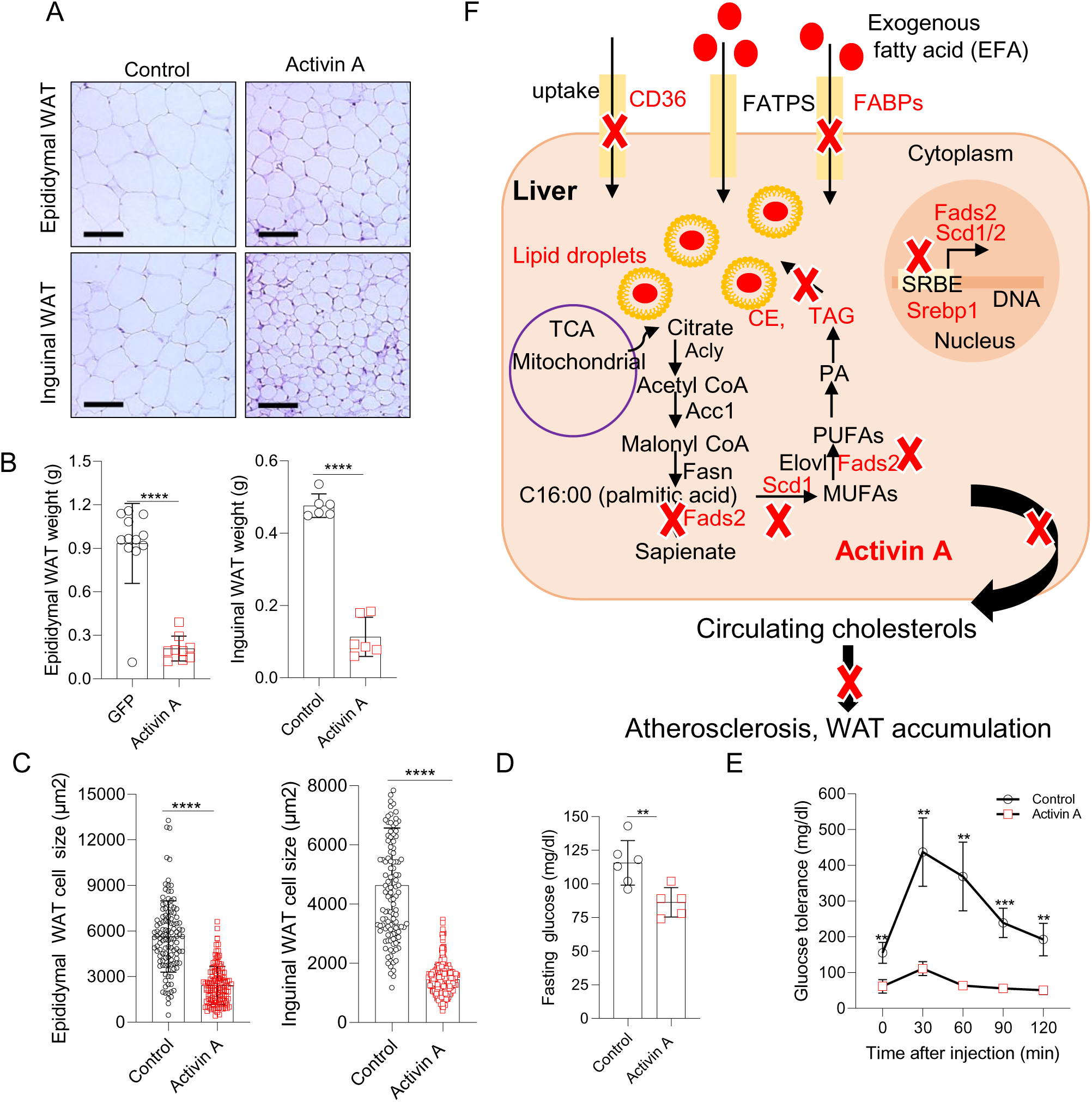
Activin A inhibits fat accumulation and improves glucose tolerance. **A.** Representative Hematoxylin and Eosin staining of epididymal White Fat Tissue (WAT) and inguinal WAT in Activin A treated mice compared with Control. N=7 for Inguinal WAT and N=11-12 in epididymal WAT. Scale bar: 50μm. **B.** Quantification of the WAT weight in Activin A delivered mice compared with Control. Unilateral fat was calculated. N=6-12 mice. **C.** Quantification of inguinal WAT and epididymal WAT adipocyte areas. N=7 mice. **D.** Fasting glucose level was measured by a glucose meter and strips. Mice were fasted for 5 hours before the measurement. N=5-6 mice. **E.** Glucose tolerance was measured by a glucometer. Overnight-fasted mice were injected intraperitoneally with glucose (2 g per kg body weight). Blood glucose levels were measured at the basal level and at 15, 30, 60, 90 and 120 min after glucose administration using a blood glucometer. N=3-4 mice. **F**. A schematic model of the role of Activin A in regulating the metabolic processing of fatty acid in the liver, and effects on circulating cholesterol levels and atherosclerosis. Activin A not only blocks genes involved in fatty acid uptake including CD36 and FABPS, but also attenuates expression of genes involved in fatty acid *de novo* lipogenic pathways including Srebp1-Scd1/Fads2. Pathways expected to be downregulated by Activin A are indicated with a red X. Scd1/Scd2 and Fads2 expression are regulated by Srebp1 at the transcriptional level. ACLY, ATP–citrate lyase; ACC, acetyl-CoA carboxylase; FASN, fatty acid synthase; ELOVLs, elongation of very long-chain fatty acid protein; PA, phosphatidic acid; TAG, triacylglycerol; CE, cholesterol ester.

### Activin A decreased inflammatory gene expression

We next explored how Activin A protects against inflammation. In overrepresentation analysis of DEG, “cytokine-cytokine receptor interaction” ranked 1 and “chemokine signaling pathway” ranked 4 among the KEGG pathways, and “cytokine-mediated signal pathway” ranked 2, “phagocytosis” ranked 5, “myeloid cell differentiation” ranked 10, and “leukocytes migration” ranked 11 among the GO Biological Process (Figure 4B and S3A). In addition, a heatmap generated for the DEG list from those pathways revealed downregulation of many genes involved in leukocyte migration (GO) and cytokine-cytokine receptor interactions, such as monocyte/macrophage trafficking/migration chemokines CCL2 (also known as MCP1), CCL3, CCL4, CCL5, CCL6, CCL959, macrophage markers CD68, CCR2 (the receptor of CCL2) and Lgals3, monocytes marker Ly-6C2, hepatic steatosis/ NASH associated lymphocyte antigen Ly6D, monocyte/macrophage homeostasis chemokine CXCL16, neutrophil trafficking chemokine CXCL1, as well as other pro-inflammatory chemokines CXCL9, CXCL10 and CXCL11 (Figure 5A). Meanwhile, liver hematopoietic lineage markers, including CD44, CD48, Fcgr1 and Csf3r were decreased in Activin A-expressing mice compared with control (Figure 5A). Next, we confirmed by QPCR that the mRNA levels of all these genes were decreased in Activin A-expressing livers compared with controls (Figure 5B-5C). We performed flow cytometry on liver samples and found that Activin A expression leads to a ∼50% decrease in F4/80+ macrophages, ∼60% decrease in Ly-6C^high^ monocytes, >50% decrease in PMN (polymorphonuclear cells), PMO (patrolling monocytes), as well as MDM (monocytes-derived macrophages) (Figure 5D-5G). In addition, eosinophils were also decreased in Activin A-expressing livers (Figure 5E, 5G). However, MM (mature macrophages or Kupffer cells) did not significantly change (Figure 5E, 5G). These data suggest that Activin A expression decreased liver inflammation and recruitment of inflammatory monocytes/macrophages and PMNs to the liver.

### Activin A inhibits fat accumulation and improves glucose tolerance

Cardiovascular diseases, including atherosclerosis, are highly correlated with obesity and diabetes^60^, and Activin A regulated genes Scd1 may also play crucial roles in obesity and diabetes. This led us to probe whether Activin A affects WAT lipogenesis and glucose tolerance. H&E staining showed that Activin A led to ∼2-fold decrease in epididymal, and inguinal WAT (Figure 6A-6B). In addition, adipocyte size was dramatically decreased in epididymal and inguinal WAT) by hepatic Activin A expression liver compared with controls (Figure 6A, 6C). Moreover, fasting glucose levels were substantially decreased by hepatic Activin A expression compared with controls (Figure 6D). Furthermore, Activin A expression in liver resulted in improved glucose tolerance (Figure 6E). Overall, these data demonstrate that hepatic Activin A inhibited fat accumulation and improved glucose tolerance in atherosclerotic mice.

## Discussion

Here we report that Activin A exogenously expressed in the liver protects against atherosclerosis. AAV8-TBG-Activin A treatment resulted in a ∼40% decrease in plasma LDL cholesterol, ∼50% decrease in plasma total cholesterol, ∼60% decrease in atherosclerotic lesions in the aortic arch, and fewer inflammatory cells infiltrating aortas and proliferating HSC in bone marrow. Hepatic Activin A not only reduced liver steatosis by inhibiting Srebp1 and Srebp2 expression and hepatic *de novo* lipogenesis, as well as exogenous fatty acid uptake, but also alleviated liver inflammation and reduced monocytes, macrophages, and neutrophil infiltration. In addition, Activin A expression in liver also reduced white fat tissue accumulation and adipocyte size as well as improving glucose tolerance. Taken together these data suggest the reduction in hepatic Activin A observed in mice on the Western diet is maladaptive and demonstrate that hepatic Activin A expression has a host of favorable effects in this context that culminate in reduced atherosclerosis and hepatic steatosis, phenotypes that cause significant morbidity and mortality throughout the world.

Increased Activin A has previously been reported in vascular lesions and plasma of patients with atherosclerosis^19^, but whether it contributed to disease pathogenesis or played a compensatory protective role has not been firmly established. While plasma Activin A was indeed increased in hyperlipidemic mice in our study, we observed a decrease in Activin A expression in the aorta and liver of these animals. Together with the finding that increasing Activin A in the liver and circulation attenuates the development of atherosclerosis, these observations suggest that the increase in Activin A reported in patient plasma and lesions may reflect a compensatory change.

While previous studies have identified anti-atherosclerotic effects of Activin A signaling on cellular processes within the vessel wall such as foam cell formation and vascular smooth muscle cell differentiation *in vitro*, our data suggest that important primary contributors to Activin’s anti-atherosclerotic effects in the LDLR^-/-^ murine model lie outside the vessel. In line with the dramatic decrease in plasma LDL and total cholesterol observed in Activin A-treated mice, our liver RNA-seq studies were consistent with anti-atherogenic changes in liver lipid metabolism, and reduced Srebp1 and Srebp2. Subsequent validation studies confirmed downregulation of key transcription factors and enzymes involved in cholesterol and triglyceride metabolism and fatty acid uptake (Srebp1 and Srebp2, Scd1/ Scd2, Fads2, Fabp1 and CD36) (Figure 6F), and reduced liver total cholesterol and free cholesterol. The reduction in cholesterol synthesis and fatty acid uptake likely contribute to the reduced inflammation seen in both the aorta and liver. Consistently, knockout of fatty acid desaturase (Fads2) reduces cholesterol and triglycerides and protects against Western diet-induced atherosclerosis^61^. In addition, bone marrow transplantation of HSC from Fabp5 deficiency mice protects against atherosclerosis in LDLR^-/-^ mice^62^. Here, Activin A had other beneficial effects including a reduction in proliferating HSCs, reduction in WAT, and improved glucose tolerance. Some of these effects may be secondary to the observed reduction in total and LDL cholesterol but other primary mechanisms may also be contributing and will be of interest for future studies.

In broad terms, many effects of Activin A (and other GDF/BMP proteins) can be seen as toggling tissues between catabolic and anabolic states. Throughout evolution in calorie scarce environments, downregulation of Activin may have been an adaptive response that conserved energy and activated useful anabolic processes. However, in calorie rich settings, exemplified here by the Western diet, the downregulation of Activin A appears maladaptive, promoting atherogenesis and hepatic steatosis, major causes of morbidity and mortality in the modern world.

Other members of the Activin/GDF/TGFβ/BMP family have also been reported to have roles in atherosclerosis. GDF11, another ligand for ACVR receptors, protected against atherosclerosis in ApoE^-/-^ mice by ameliorating inflammation and endothelial cell injury^63^. Knockout of the TGFβ receptor, TGFβRII, in dendritic cells, smooth muscle cells and T cell was reported to accelerate atherosclerosis while knockout of TGFβRII in endothelial cells protected against atherosclerosis^64-67^. Neutralization of TGFβ with an antibody (anti–hTGF-β1, -β2, -β3 2G7 monoclonal) increased atherosclerosis and inflammation^68^. Interestingly, TGFβ1 was also reported to induce HSC quiescence and cell cycle arrest by upregulation of p57, a member of the cyclin-dependent kinase inhibitor family69, 70. Bone marrow transplantation studies showed that the hematopoietic stem/progenitor population was reduced in bone marrow from TGFβ1KO pups and bone marrow reconstitution was impaired compared with TGFβ1+/+ pups^71^. However, TGFβ1 receptor I (TGFβRI) deficient mice showed normal HSC self-renewal and regeneration ability^72^. This is partially because TGFβ2, another isoform of TGFβ, supports LSK cell proliferation^73^. It will be interesting to probe the role of Activin A, its receptors, as well as other ligands in Activin pathway in HSC self-renewal, regeneration, expansion, and atherosclerosis.

Interestingly, we did not see pro-fibrotic or pro-apoptotic effects of hepatic Activin A expressions despite prior reports of such effects with other members of this protein family. Rather, we instead saw a robust decrease in fibrosis markers and no alteration in apoptosis in the liver (data not shown) in hyperlipidemic mice treated with Activin A. Unlike the classic TGFβ signaling pathways, Activin A is regulated by a negative feedback loop. FST is not only a direct target but also a strong antagonist of these pathway. In addition, Activin A also forms a non-signaling complex with the type I receptors (ACVR1a, ALK2) and type II receptor. These self-limiting mechanisms may preserve homeostasis and prevent overactivation of the Activin A pathway. The expression level of Activin A is likely a key determinant of whether these self-limiting pathways are activated or not, with beneficial or harmful consequences for the liver. Here we delivered a very low dosage of AAV8-TBG-Activin A (2×10^10 genome copies/mice, ∼2000 pg/ml in plasma, compared to the common dose of 1-2×10^11 genome copies/mouse^32-34^). This level of expression activated downstream signaling, as indicated by an increase in phosphorylation of Smad2/3, as well as increased FST. The Activin A receptors ACVR1b, ACVR2a, and ACVR2b were also decreased or showed a nonsignificant trend toward decrease. Interestingly, levels of the liver injury marker plasma ALT greatly decreased in Activin A expressing liver compared with control, suggesting a low dosage of AAV8-TBG-Activin A may be beneficial for the liver. This is consistent with a previous study showing low-dose AAV2/8-smad3 (2×10^9 genome copies/mice) attenuated aortic atherosclerosis without increasing fibrosis^74^.

The beneficial effects of hepatic Activin A expression raise the intriguing possibility that Activin A expression could have therapeutic potential. However, reasonable concerns could be raised about this approach. We have previously shown that Activin A levels increase with age in humans and contribute to multiple models of cardiac dysfunction and heart failure^75^. Activin A was sufficient to cause cardiac dysfunction in healthy mice but importantly, that was at higher concentrations of Activin A (∼10,000 pg/ml) by adenovirus delivery without special diet^75^. Here, the levels of Activin expression used were lower and echocardiography did not reveal alterations in cardiac function (data not shown). Still there could be legitimate concerns about therapeutic windows particularly given the challenges of fine-tuning concentrations achieved with gene therapy approaches. Finally, while the growing number of treatments available for primary and secondary prevention of atherosclerosis may undermine interest in that setting, no drugs have been approached for hepatic steatosis or the closely related non-alcoholic liver disease suggesting there may be greater enthusiasm for evaluation of Activin A’s ability to address this important unmet clinical need.

Activin A also reduced WAT and improved glucose tolerance, suggesting a potential beneficial role for Activin A in obesity and diabetes, similar with hepatic Activin E^76^. However, this may seem paradoxical given that *deletion* of a closely related protein, myostatin (GDF8) has been shown to reduce obesity and protect against diabetes in mice^77^. It seems possible that deletion and overexpression could culminate in similar phenotypes through different mechanisms. For example, the dramatic skeletal muscle growth seen in myostatin knockout mice likely mediates multiple metabolic benefits while the catabolic state induced by overexpression of Activin A may reduce fat storage and insulin resistance. In this context, it would be interesting in future studies to examine the effects of genetic or pharmacological inhibition of these pathways in atherosclerosis as well.

In summary, the data presented here reveal for the first time that Activin A expressed in the liver protects against atherosclerosis, as well as liver steatosis and inflammation, reduces fat accumulation and improves glucose tolerance. These findings have implications for our understanding of the role of this pathway and the regulation of processes central to atherogenesis, while suggesting the therapeutic potential of moderate hepatic Activin A expression in atherosclerotic vascular disease and hepatic disorders, such as obesity and non-alcoholic liver disease, warrants further investigation.

## Non-standard Abbreviations and Acronym

TBG: thyroxine binding globulin
CVD: cardiovascular diseases
LDL: low density lipoprotein
VLDL: very low-density lipoprotein
oxLDL: (oxidized-LDL)
AAV: adeno-associated virus
FST: Follistatin
HSC: hematopoietic stem cell
LtHSC: long-term hematopoietic stem cell
StHSC: short-term hematopoietic stem cell
MPP: multipotential progenitor cells
PMN: polymorphonuclear cells
DEG: differentially expressed genes
FAs: fatty acids

## Acknowledgments

We acknowledge Suying Liu (Children’s Hospital of Philadelphia), Dr. Shun He and Dr. Alexandre Paccalet (Center for Systems Biology, MGH), Dr. Anlu Chen (Broad Institutes) for their technical supports and professional suggestions, Danian Cao (Wellman Center Photopathology Core, MGH) for flow cytometry analysis, Yoshiko Iwamoto (Center for Systems Biology, MGH) for tissue staining, and Photopathology Core Wellman Center, MGH CCM Clinic Pathology Lab, and MGH Histopathology Research Core for their services.

H.L, and A.R conceived this study. H.L designed and performed experiments, analyzed data, wrote, and edited the manuscript. M.H.H wrote and edited the manuscript. C.Y.X performed echocardiography. R.K. J.B.G, H.L, and A.K analyzed RNA-seq data. A.R supervised the study, analyzed data, wrote, and edited the manuscript, and secured funding.

## Supplemental Material

Supplemental Materials and Methods

Supplemental Table 1

Supplemental Figure S1, S2, and S3

Supplemental References 29-31

---

**Figure S1, related to Figure 1.**
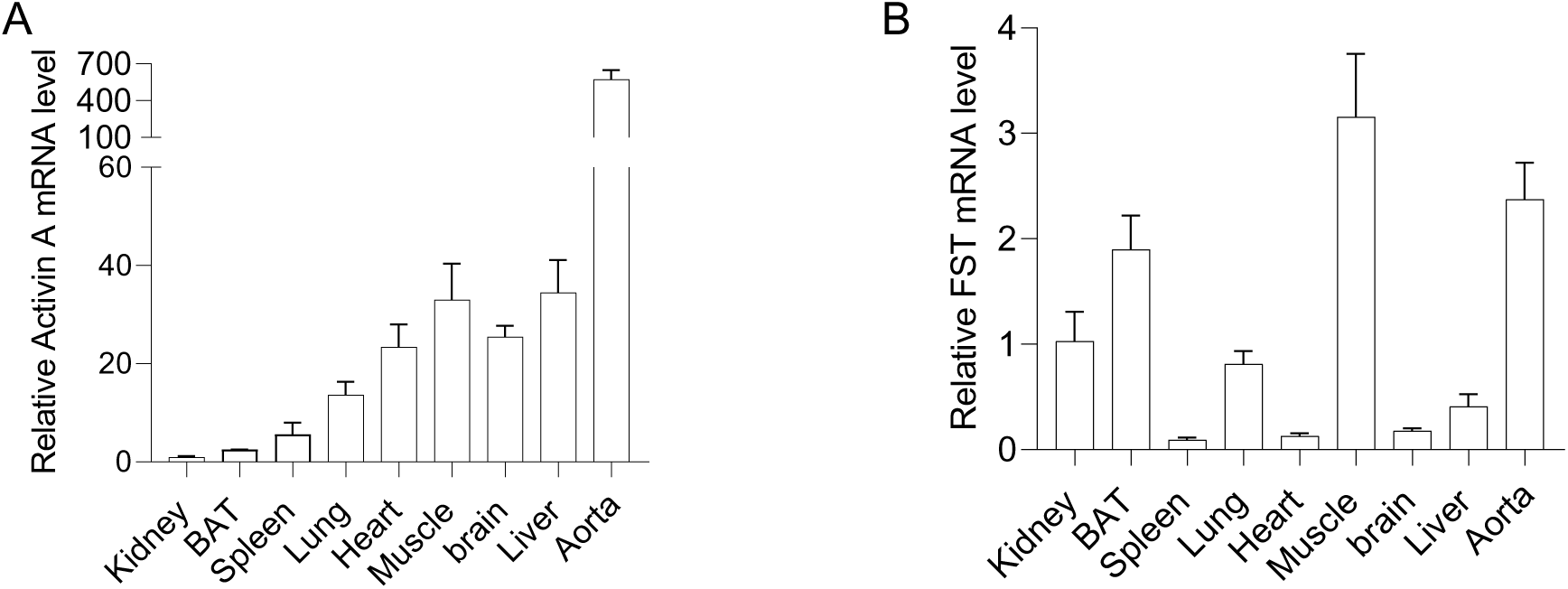
**A-B**. mRNA levels of Activin A and FST measured by QPCR in different tissues in mice.

**Figure S2, related to Figure 2.**
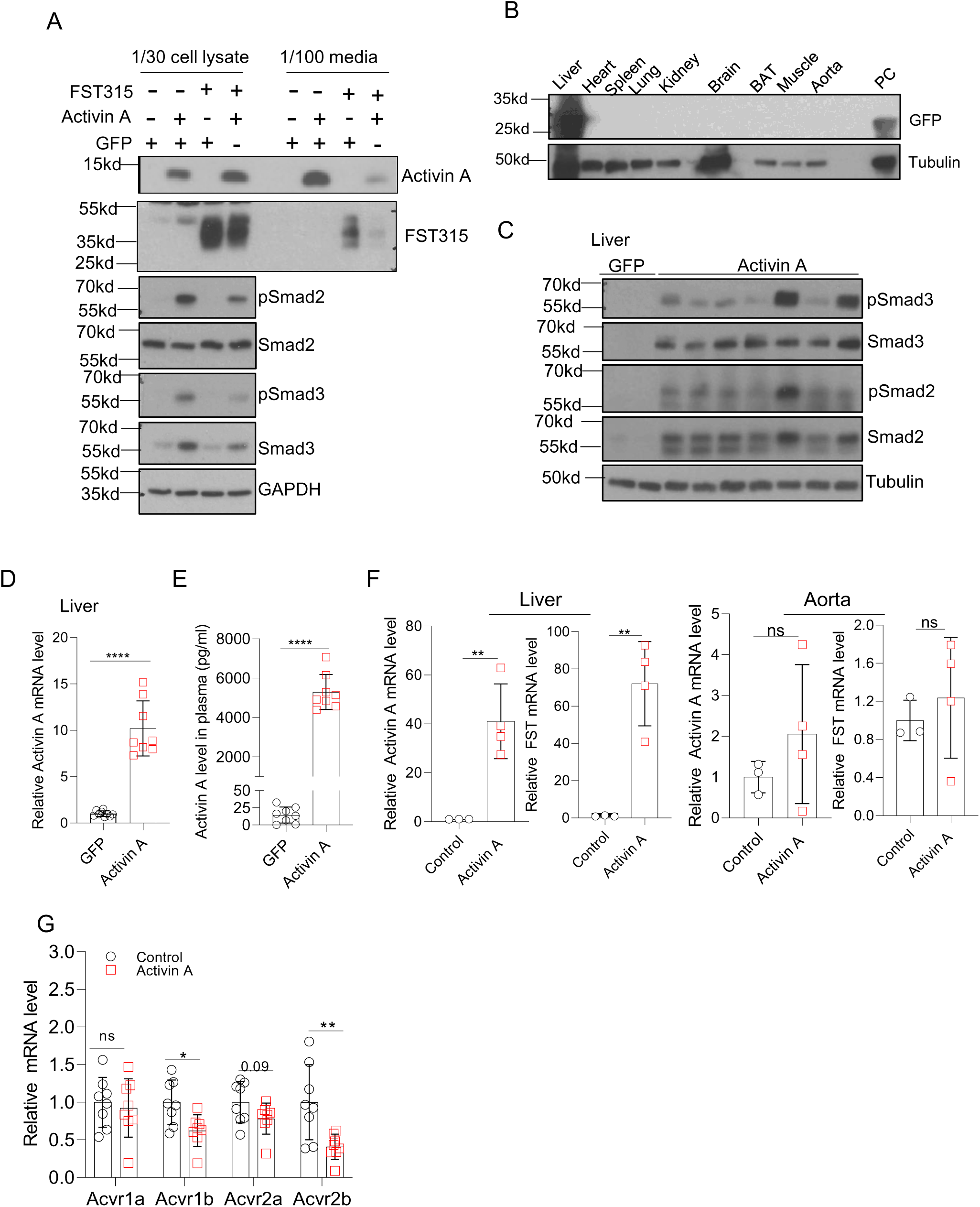
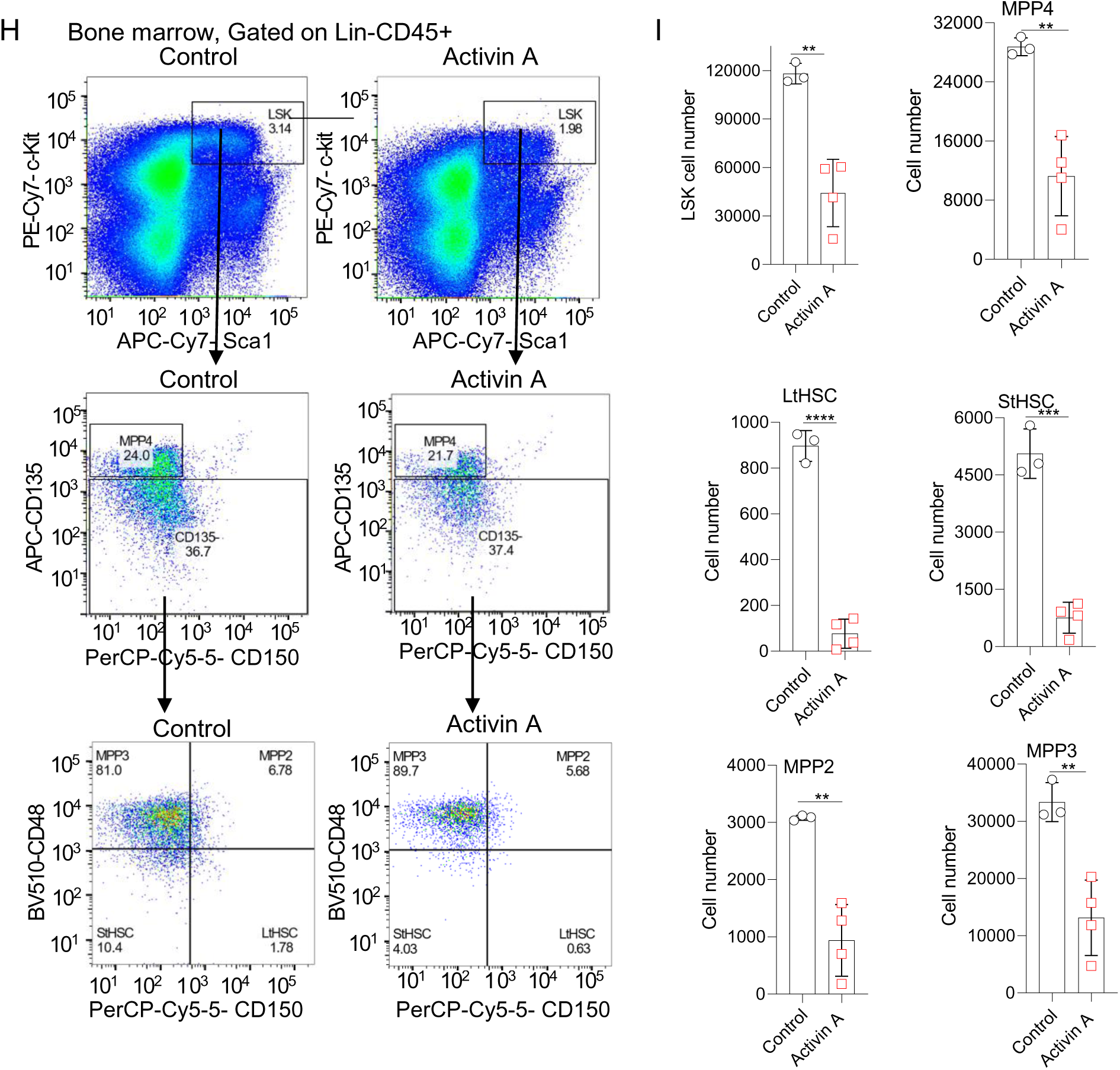
**A.** Western blotting showing that AAV8-TBG-Activin A is expressed in HepG2 cells, activates Smad2/Smad3 phosphorylation, and circulates in the media, while AAV8-TBG- FST315 blocks the activity of Activin A. The indicated plasmids were transfected into HepG2 cells and cultured for 48 h. **B.** Western blotting shows AAV8-TBG-GFP was highly expressed in liver but no other tissue after tail vein injection. 6-week-old male C57/BL6 mice were I.V. injected with 5×10^10 AAV8- TBG-GFP, and then sacrificed after 4 weeks. Protein levels were quantified and normalized by BCA kit (pierce), and target proteins were detected by western blotting using anti-GFP antibody. HepG2 cell lysates with GFP expression were used as the positive control (PC). 18.74 μg protein /sample. **C-D**. Western blotting and Q-PCR show AAV8-TBG-Activin A was expressed and activated the Activin pathway. 6-week-old male C57/BL6 mice were I.V. injected with 5×10^10 genome copies AAV8-TBG-GFP, and then scarified after 4 weeks. Protein level was quantified by BCA kit (pierce) and target proteins detected by western blotting. **E**. ELISA shows liver expressed Activin A circulates in plasma. Mice from H were used. Plasma FST levels are close to baseline (0) and not shown here. **F.** Q-PCR shows AAV8-TBG-Activin A is expressed and induces expression of its direct target gene FST in the liver but not aorta in atherosclerotic LDLR^-/-^ mice. Control, n=3 and Activin A, n=4. **G.** Q-PCR shows the mRNA levels of Activin pathway receptors in control and Activin A-expressing atherosclerotic liver. N=8 mice. Acvr1a, Acvr1b, Acvr1c, Acvr2a and Acvr2b primers were used for Q-PCR, and Acvr1c was undetectable. **H**. Assessment and gating strategy of hematopoietic stem cells (HSC) populations by flow cytometry. LSK: Lin2-CD45+Sca1+c-Kit+. Long-term HSC (LtHSC), short-term HSC (StHSC) and multipotent progenitors (MPP2, MPP3 and MPP4). StHSC (CD135-/CD150-/CD48- LSK), LtHSC (CD135-/CD150+/CD48- LSK) MPP2 (CD135-/CD150+/CD48+ LSK), MPP3 (CD135- /CD150-/CD48+ LSK), and MPP4(CD135+/CD150-/CD48+/- LSK). **I.** Quantification of the number of LSK, LtHSC, StHSC and multipotent progenitors in control and Activin A-expressing atherosclerotic mice. N=3-4 mice. Similar results were obtained from 3 independent experiments.

**Figure S3, related to Figure 4.**
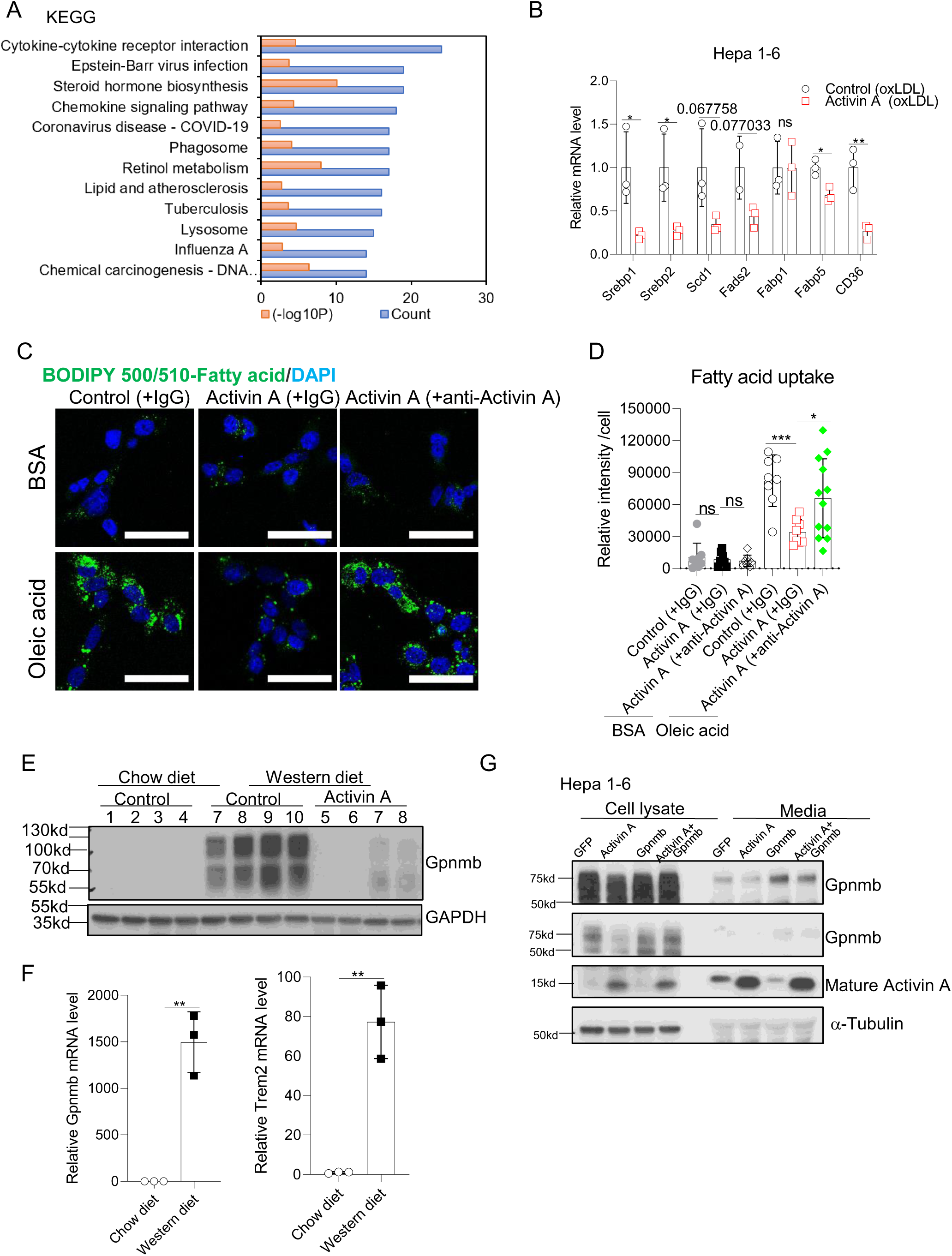
**A.** KEGG enrichment of top 12 significantly changed pathways in RNA-seq analysis of Activin A-expressing liver compared with control. TPM (control) or TPM (Activin A) ≥ 5, p<0.05, |log2Fc|≥ 1. **B.** Q-PCR shows the mRNA levels of genes for hepatic *de novo* lipogenesis and exogenous fatty acid uptake decrease in Hepa 1-6 cells treated with Activin A-expressing conditioned media and 10 μg/ml oxLDL compared with control. Hepa 1-6 cells were transfected with TBG-GFP or TBG-Activin A plasmid, and 24h later were changed into 0.5%BSA (fatty acid free)/RPMI at 37°C for 24h. The conditioned media were collected and filtered through a 0.22 μm filter, diluted twice with fresh 0.5%BSA/RPMI1640, and then added into fresh Hepa 1-6 cells together with 10 μg/ml oxLDL for 24 hours. **C-D.** Confocal images and quantification of fatty acid uptake. Hepa 1-6 cells were transfected with TBG-GFP or TBG-Activin A plasmid, and 24h later were changed into 0.5%BSA (fatty acid free)/RPMI at 37°C for 24h. The conditioned media were collected and filtered with a 0.22 μm filter, diluted twice with fresh 0.5%BSA/RPMI1640, and then added into fresh Hepa 1-6 cells, with or without 30 μΜ Oleic acid (OA), with 1μg/mL Mouse IgG1 Isotype Control or neutralizing anti-Activin A antibody for 24 hours. Then cells were washed with PBS, stained with 2 μΜ BODIPY 500/510-fatty acid, fixed with 4%PFA, and stained with DAPI. Scale bar, 50 μm. **E.** Western blotting shows Gpnmb is expressed in western diet-fed LDLR^-/-^ control mice compared with chow but decreases in Activin A-expressing western diet-fed LDLR^-/-^ mice. **F.** Q-PCR shows Gpnmb and Trem2 mRNA increase in western diet-fed LDLR^-/-^ mice compared with chow diet. **G.** Western blotting shows Activin A decreases Gpnmb protein levels in cell lysate and media. α-Tubulin as the loading control.

