## supplemental methods and supplemental table1 for "Beneficial Effects of Moderate Hepatic Activin A Expression on Metabolic pathways, Inflammation, and Atherosclerosis"

*Short title: Beneficial Effects of Activin A in Atherosclerosis*

*Total Word Count:2078*

Cardiovascular Research Center, Massachusetts General Hospital, and Harvard Medical School, Boston, MA, 02114, USA.

**Supplementary Materials includes:**

Supplemental Materials and Methods

Supplemental Table 1

Supplemental Figure S1, S2, and S3

Supplemental References 29-31

### Supplemental Materials and Methods

#### Molecular Cloning

The AAV8-TBG-GFP plasmid was generated by replacing the Ctnt promoter in the pENN.AAV.Ctnt plasmid (Addgene 105543) with the liver-specific TBG promoter[27] using ApaI and BstAPI digestion. To produce the AAV8-TBG-Activin A plasmid, GFP was then replaced with Activin A coding sequence from Origene (#MC206397) using NcoI and KpnI.

#### AAV Production, Purification, and Titration

AAV production, purification and titration were performed as described[28, 29]. Briefly, HEK293 cells were transfected with pΔF6 (Addgene Plasmid #112867), pAAV2/8 (Plasmid #112864) helper plasmids and AAV8-TBG-GFP or AAV8-TBG-Activin A transfer plasmids by PEI. Viruses were purified using iodixanol gradient ultracentrifugation using OptiPrep™ Density Gradient Medium (Sigma D1556), concentrated with columns（Fisher scientific UFC910024）, and titrated using droplet digital PCR.

#### OxLDL Uptake

Fresh peritoneal macrophages were incubated with 4μg/ml Dil-oxLDL (Thermo Fisher, #L34358) in 5% BSA (fatty acid-free)/RPMI1640 at 37°C for 2hrs, then fixed with 4%PFA for 15min at room temperature, permeabilized with 0.1%Triton X-100 for 5min, washed and mounted with ProLong® Gold Antifade Mounting with DAPI (Thermo Fisher Scientific), imaged by confocal microscope (Leica sp8), and quantified with Image J software.

##### Fatty acid uptake

Hepa 1-6 cells (ATCC) were transfected with TBG-GFP or TBG-Activin A plasmid by Lipofectamine 3000 (Thermo Fisher Scientific) for 24h, then changed into 0.5%BSA (fatty acid free)/RPMI 1640 at 37°C for 24h. Conditioned media were collected and filtered (0.22 μm), diluted twice with fresh 0.5%BSA (fatty acid-free)/RPMI1640, and added to fresh Hepa 1-6 cells, with or without 30μM Oleic acid (OA, Sigma) with 1μg/mL Mouse IgG1 Isotype Control (R&D, MAB002) or neutralizing anti-Activin A antibody (R&D, MAB3381) for 24 hours. Cells were washed 3 times with PBS, stained with 2 μM BODIPY 500/510-fatty acid (Thermo Fisher Scientific, D3823) for 30min, fixed with 4%PFA at room temperature for 15min, permeabilized with 0.1%Triton X-100 for 5min, and mounted with ProLong® Gold Antifade Mounting with DAPI (Thermo Fisher Scientific), imaged by confocal microscopy (Leica sp8), and quantified with Image J software.

##### Glucose Tolerance Test

Overnight-fasted mice were injected intraperitoneally with glucose (2gm/kg body weight). Blood glucose levels were measured at baseline and 15, 30, 60, 90 and 120min after glucose administration using a blood glucometer (Clarity BG1000).

##### Blood and Plasma Analyses

Fasting blood glucose was measured using a glucometer (Clarity BG1000 Blood Glucose Meter and Strips, Clarity, CD-BG5) in mice after fasting for 5hr with free access to water. Overnight fasting plasma was collected in 5 mM EDTA. Plasma ALT (SGPT) and lipids including total cholesterol, total triglyceride, HDL, and LDL were analyzed by IDEXX BioAnalytics. Plasma Activin A and FST levels were quantified by Human/Mouse/Rat Activin A Quantikine ELISA Kit (R&D, DAC00B) and Human Follistatin Quantikine ELISA Kit (R&D, DFN00).

#### Quantitative RT-PCR

Total RNA was extracted from liver tissue using Trizol (Thermo Fisher Scientific)/chloroform/isopropyl alcohol. cDNA was synthesized by reverse transcription using PrimeScript RT Master Mix (Takara) and subjected to real-time PCR (Thermo Fisher Scientific, QuantStudio™ 6 Flex Real-Time PCR System, 384-well) with gene-specific primers in the presence of SYBR Green (Applied Biosystems). Relative abundance of mRNA was calculated by normalization to hypoxanthine phosphoribosyltransferase 1 (HPRT) mRNA. For primers used see Supplemental Table 1.

#### Immunoblotting

Liver protein was extracted using RIPA/SDS buffer (Thermo Fisher Scientific, #89900), and protein concentration was quantified by a BCA assay (Thermo Fisher Scientific, #23227). Proteins were resolved by electrophoresis on 4-20% Tris gels (Bio-Rad) and transferred to PVDF membranes (Millipore Sigma). Membranes were blocked for 1hr at room temperature in Tris-buffered saline and 0.1% Tween 20 (TBST) containing 5% (wt/vol) nonfat milk and incubated with primary antibodies in the same buffer at 4°C overnight, using 1:1000 dilution. Protein bands were detected with horse radish peroxidase-conjugated secondary antibodies. Primary antibodies used for immunoblotting: anti-Activin A (R&D, AF338, only for exogenous western blotting *in vitro*), anti-FST (R&D, AF669), anti-Smad2 (Cell Signaling Technology, 5339S), anti-Smad3 (Cell Signaling Technology, 9513S), pSmad2 (Ser465/467, Cell Signaling Technology, 3108S) pSmad3 (Ser 423/Ser 425, Abcam, ab52903), Srebp1(Santa Cruz, sc-13551), Srebp2 (Abcam, ab30682), anti-Gpnmb (R&D, AF2330), anti-CD36 (R&D, AF2519), anti-Fabp1 (Cell Signaling Technology, 13368T), anti-Fabp5 (R&D, AF1476). The protein signals were detected by ECL Detection Reagent (Thermo Fisher Scientific) and quantification by Image Studio software.

#### Atherosclerotic Lesion Analysis

For *en face* analyses, the aorta was removed from the heart to 5-10mm after bifurcation of the iliac arteries and including the subclavian right and left common carotid arteries, stained with oil red, and imaged (Olympus SZX16 microscope). Total atherosclerotic plaque area was quantified using ImageJ software.

#### Liver and Fat Histology

Hepatic and fat tissue were fixed in neutral buffered 10% formalin (Sigma) for 48 h at room temperature, embedded and processed in the pathology core according to standard protocols. Briefly, formalin fixed tissues were processed for dehydration with gradient ethanol solutions (70%, 95%, 100%) and cleared with 100% CitriSolv (xylene replacement). Tissues were put in hot paraffin liquid (60°C) for 1hr for 3 times. Routine Hematoxylin and Eosin (H&E) staining was performed on 5μm paraffin sections using a Leica ST5010 Autostainer XL. For immunohistochemistry (IHC) staining, paraffin sections were deparaffinized and rehydrated through graded ethanol solutions, immersed in antigen retrieval citrate buffer (Sigma, C9999) at 95°C for 30min. Sections were blocked with 10% goat serum, stained with antibodies indicated using the avidin–biotin complex system (Vector Laboratories). 3,3′-diaminobenzidine (DAB) was used as the substrate. Nuclei were counterstained with hematoxylin. Collagen was stained with Picrosirius (Sirius) Red (Polysciences, # 24901) according to instructions. For frozen sections, mouse livers were fixed in 4% PFA at 4°C for 48hr, rinsed with PBS, and incubated with 30% sucrose for 12–16 hours at 4°C, then embedded in Tissue-Tek OCT compound (Sakura Finetek). Lipid was stained by oil red on frozen sections. Slides were scanned using a NanoZoomer 2.0-RS (Hamamatsu, Japan) and quantified. Frozen sections were subjected to immunofluorescent staining with anti-CD45 (BD, 550539), anti-F4/80 (Abcam, ab6640), anti-Gpnmb (R&D, AF2330) anti-HNF4α (R&D, PPH141500), followed by the secondary antibodies Alexa Fluro 488 Rabbit-anti-Rat (Thermo Fisher Scientific, A-21210 ), Rabbit-anti-goat 568 (Thermo Fisher Scientific, A11079), Rabbit-anti-mouse 647 (Thermo Fisher Scientific, A-21239 ), and DAPI (nuclei), imaged by confocal microscope (Leica sp8), and then quantified by Image J software.

**Tissue Cholesterol Analysis**

For liver cholesterol quantification, liver tissue (20mg) was washed in cold PBS, homogenized in 200μL diluted NP40 lysis buffer with protease inhibitors ( from Triglyceride Colorimetric Assay Kit, Cayman 10010303), and centrifuged at 10,000×g for 10min. Supernatants were transferred to Eppendorf tubes, mixed thoroughly, and triglyceride (Triglyceride Colorimetric Assay Kit, Cayman 10010303), cholesterol (Wako, #439-17501), and free cholesterol (Wako, #994-02501) were measured according to the manufacturer's instructions.

**BrdU incorporation**

BrdU incorporation was performed as described[30]. Briefly, 1mg BrdU (BD Biosciences) was intraperitoneally injected 2 hr before euthanasia. A BrdU flow kit (BD Biosciences) was used to stain BrdU+ cells.

**Flow Cytometry**

Flow cytometry was performed as described[30, 31]. Aortae and livers were excised after PBS (Thermo Fisher Scientific) perfusion, minced, and digested with 450Uml/L collagenase I (Worthington Biochemical), 125Uml/L collagenase XI, 60Uml/L DNase I and 60Uml/L hyaluronidase (Sigma-Aldrich) in PBS at 37°C (Aortae for 40min and livers for 20 min). Bone marrow cells were collected from mouse femur and tibia by flushing bones with PBS and a single-cell suspension was made by passing cells through a 27-gauge needle. Aortae, livers, and bone marrow cells were passed through a 70μm-filter, and red blood cells were lysed with RBC lysis buffer. Total viable cell numbers were counted using counting beads (Thermo Fisher Scientific). Single-cell suspensions were stained in PBS supplemented with 2% FBS and 0.5% BSA. DAPI or Zombie Aqua™ Fixable Viability Kit were used for staining live/dead cells. Monoclonal antibodies used for flow cytometry are listed below:

Fc blocking: anti-CD16/32 antibody (clone 2.4G2, TONBO Biosciences, #70-0161-U500). Anti-CD45 (BioLegend, clone30-F11, 103108), anti-mouse CD45.2 (BioLegend, clone 104, 109807), anti-mouse Ly-6C (Biolegend, clone HK1.4, 128011), anti-mouse Ly-6G (Biolegend, clone 1A8, 127623), anti-F4/80 Rat Monoclonal Antibody (Biolegend, clone BM8, 123113 and 123115), anti-MHCII (eBioscience, clone M5/114.15.2, 17-5321-81), anti-CD11b (BioLegend, clone M1/70, 101245 and 101215), anti-CD127 (IL-7Ra) ( BioLegend, cloneSB/199, 121111), anti-TER-119 (BioLegend, clone TER119, 116207), anti-NK1.1 ( BioLegend, clone PK136, 108707), anti-CD19 (BioLegend, clone 6D5, 115507), anti-CD90.2 ( BioLegend, clone 30-H12, 105307), anti-CD3 (Biolegend, clone 17A2, 100205), PE anti-mouse Lineage Cocktail with Isotype Ctrl and FITC anti-mouse Lineage Cocktail with Isotype Ctrl (Biolegend, 133301 and 133303, including anti- CD3ε, clone 145-2C11; anti- Ly-6G/Ly-6C, clone RB6-8C5; anti- CD11b, clone M1/70; anti- CD45R/B220, clone RA3-6B2; anti-TER-119/Erythroid cells, clone Ter-119), anti-CD150 (Biolegend, clone: TC15-12F12.2, 115921), anti-CD135 (Biolegend, cloneA2F10, 135309), anti-CD117 (c-Kit)( Biolegend, clone 2B8, 105813), anti-Ly-6A/E (Sca-1)(Biolegend, clone D7 108125), anti-mouse CD48 (Biolegend, clone HM48-1, 103443), anti-Brdu (eBioscience, clone BU20A, 17-5071-41), anti-Trem2 (Thermo Fisher Scientific, clone 78.18, MA5-28223), and anti-CD9 (Biolegend, clone MZ3, 124805). Lineages were defined as: Lin1: Ter119/NK1.1/CD19/CD90.2/CD3, and Lin2: CD3/Ly-6G/Ly-6C/B220/CD11b/Ter119/ CD127.

For aortae: Ly-6C$^{high}$ monocytes (CD45+Lin1−CD11b+F4/80−Ly-6C$^{high}$), macrophages (CD45+Lin1−CD11b+F4/80+Ly-6C$^{low}$), and neutrophils (CD45+Lin1−CD11b+Ly-6G+F4/80−). For bone marrow: LSK cells (CD45+Lin2−c-Kit+Sca1+), multipotent progenitors MPP2 (CD135-/CD150+/CD48+LSK), MPP3 (CD135-/CD150-/CD48+LSK), and MPP4 (CD135+/CD150-/CD48+/- LSK), short-term hematopoietic stem cells (StHSC, CD135-/CD150-/CD48- LSK), long-term hematopoietic stem cells (LtHSC: CD135-/CD150+/CD48- LSK). For liver as described[31]: PMN (polymorphonuclear cells, CD45+CD11b+Ly-6G+, including neutrophils, basophils, and mast cells), PMO (patrolling monocytes, CD45+CD11b+Ly-6G-Ly-6C- MHC-II-), MDM (monocytes-derived macrophages, CD45+CD11b+Ly-6G-Ly-6C$^{high}$ MHC-II$^{high}$), MM (mature macrophages, also known as Kupffer cells, CD45+CD11b+Ly-6G-Ly-6C$^{neg/low}$ MHC-II $^{high}$), ED (eosinophils, CD45+CD11b+Ly-6G-Ly-6C$^{low}$ MHCII-), Ly-6C$^{high}$ monocytes (CD45+CD11b+Ly-6G-Ly-6C$^{high}$ MHCII-).

**RNA-seq and Bioinformatic Analyses**

Total RNA was isolated from liver using Trizol (Thermo Fisher Scientific)/chloroform/isopropyl alcohol. RNA quality was analyzed using an Agilent Tape Station with all RNA Integrity Numbers (RIN) >8.5. Sequencing libraries were generated from total RNA (1μg) using the TruSeq stranded mRNA kit (Illumina Corp.) and sequenced using NovaSeq6000 (20M PE150 Read Pairs) by Novogne company (USA).

Prior to gene expression analysis, Fastqc was used to determine the sequence quality after adapters and low-quality reads were trimmed with Trim-galore. Reads were aligned to the C57BL/6J mouse genome by Hisat2 using the GRCm38 (mm10) reference sequence. Sequence alignment files were written into bam files, sorted, and indexed by Samtools (version 1.9.0) and genomic reads were counted by HTSeq (v0.11.1). Differential gene expression was analyzed and normalized with R package DESeq2 and limma. Statistical significance was calculated using Wald test. R package ClusterProfiler (version 3.8) was applied to rank KEGG and Go enrichment based on the shrunken limma log2 fold changes. Genes with an average TPM (Transcripts Per Million) $\geq 5$ in $\geq 1$sample, adjusted p-value$<0.05$ and absolute log2 (fold change) $\geq 1$ were included. The heatmap of the top KEGG pathway after a cutoff of adjusted p-value$<0.05$, TPM$\geq 5$, absolute log (fold-change) $\geq 2$ was generated using R package heatmap (v.1.0.12) on Z-scale normalized counts by log2 (TPM+0.05).

#### Supplemental Table 1. List of primers used for Q-PCR

| primer name | primer sequence |
|---|---|
| Activin A-RT-F | ACGACATTGGCAGGAGG |
| Activin A-RT-R | GCTGAAATAGACGGATGGTG |
| Col1a1-RT- F | GCTCCTCTTAGGGGCCACT |
| Col1a1-RT- R | CCACGTCTCACCATTGGGG |
| Col1a2-RT- F | GTAACTTCGTGCCTAGCAACA |
| Col1a2-RT- R | CCTTTGTCAGAATACTGAGCAGC |
| Col3a1-RT-F | CTGTAACATGGAAACTGGGGAAA |
| Col3a1-RT- R | CCATAGCTGAACTGAAAACCACC |
| Srebp2-RT-F | CCAAAGAAGGAGAGAGGCGG |
| Srebp2-RT-R | CGCCAGACTTGTGCATCTTG |
| Srebp1-RT-F | CAAGGCCATCGACTACATCCG |
| Srebp1-RT-R | CACCACTTCGGGTTTCATGC |
| FST-RT-F | TGCTGCTACTCTGCCAGTT |
| FST-RT-R | CAACACTCTTCCTTGCTCAGT |
| ACVR1a-RT-F | GGCTGCTTTCAGGTTTATGA |
| ACVR1a-RT-R | GGACTTCCCTTTAGTGGGC |
| ACVR1b-RT-F | CGTCTTCCTGGTCATCAACTAT |
| ACVR1b-RT-R | GCGTCTTGTCTTTGGAGAGA |
| ACVR1c-RT-F | CAAGGTAAGCCTGCTATTGC |
| ACVR1c-RT-R | GGTTCCCACTTTAGGATTCTG |
| Acta2-RT- F | ATGCTCCCAGGGCTGTTTTCCCAT |
| Acta2-RT- R | GTGGTGCCAGATCTTTTCCATGTCG |
| Tnfα-RT- F | CTTCTGTCTACTGAACTTCGGG |
| Tnfα-RT-R | CAGGCTTGTCACTCGAATTTTG |
| Mcp1-RT-F | TTAAAAACCTGGATCGGAACCAA |
| Mcp1-RT-R | GCATTAGCTTCAGATTTACGGGT |
| CCL2-RT-F | GCATCCACGTGTTGGCTCA |
| CCL2-RT-R | CTCCAGCCTACTCATTGGGATCA |
| CCL2-RT-F | AAGGAATGGGTCCAGACA |
| CCL2-RT-R | GTGCTTGAGGTGGTTGTG |
| CCL3-RT-F | TGTACCATGACACTCTGCAAC |
| CCL3-RT-R | CAACGATGAATTGGCGTGGAA |
| CCL4-RT-F | AAACCTAACCCCGAGCAACA |
| CCL4-RT-R | CCATTGGTGCTGAGAACCCT |
| CCL5-RT-F | TGCCCACGTCAAGGAGTATTT |
| CCL5-RT-R | TCCTAGCTCATCTCCAAATAGTTGAT |
| Ccl6-RT-F | AAGAAGATCGTCGCTATAACCCT |
| Ccl6-RT-R | GCTTAGGCACCTCTGAACTCTC |
| CCL9-RT-F | CCCTCTCCTTCCTCATTCTTACA |
| CCL9-RT-R | AGTCTTGAAAGCCCATGTGAAA |
| CXCL1-RT-F | GGCTGGGATTCACCTCAA |
| CXCL1-RT-R | GCTTCAGGGTCAAGGCAA |
| CXCL2-RT-F | GCGCTGTCAATGCCTGAAGA |
| CXCL2-RT-R | TTTGACCGCCCTTGAGAGTG |
| CXCL9-RT-F | TCCTCTTGGGCATCATCTTCC |
| CXCL9-RT-R | TTTGTAGTGGATCGTGCCTCG |

| | |
|---|---|
| CXCL10-RT-F | TGCTGGGTCTGAGTGGGACT |
| CXCL10-RT-R | AGGATAGGCTCGCAGGGATG |
| CXCL11-RT-F | ACAGGAAGGTCACAGCCA |
| CXCL11-RT-R | ACTTTGTCGCAGCCGTTA |
| CXCL14-RT-F | ACCCACACTGCGAGGAGAA |
| CXCL14-RT-R | AAGCGTTTGGTGCTCTGC |
| CXCL14-RT-F | ATCGTCACCACCAAGAGC |
| CXCL14-RT-R | TCTCGTTCCAGGCATTGT |
| CXCL16-RT-F | ACATTTGCCTCAAGCCAG |
| CXCL16-RT-R | AGGGTTGGGTGTGCTCTT |
| CD68-RT-F | ACCACAAATGGCACTGCT |
| CD68-RT-R | CTGAACACAAGGCTGGGA |
| ly6C2-RT-F | CGCCTCTGATGGATTCTG |
| ly6C2-RT-R | TCCTTGATTGGCACACCA |
| ly6C2-RT-F | CAAAGAAGGAAACTAAAGACCC |
| ly6C2-RT-R | AGGCTGAACAGAAGCACC |
| Ly6D-RT-F | ATGCAACGAGAGGCTGGTCA |
| Ly6D-RT-R | ACTAGAAGGGAAGGGGCAAG |
| Ly6e-RT-F | CACTACTGGGCCTTGGACTC |
| Ly6e-RT-R | CTGAAGGTTCCTAGGGTCCC |
| TLR1-RT-F | TCAGCACTACGATCGGTTTG |
| TLR1-RT-R | TGTTTCCACATTGTTCAGGG |
| TLR2-RT-F | GCGACATCCATCACCTG |
| TLR2-RT-R | TCATCTACGGGCAGTGG |
| Fabp5-RT-F | GCCTGATGGAAAGCCAC |
| Fabp5-RT-R | CCGTGATGTTGTTGCCA |
| Scd1-RT-F | GCAAGCTCTACACCTGCCTCTT |
| Scd1-RT-R | CGTGCCTTGTAAGTTCTGTGGC |
| Cd36-RT-F | CGTGGCAAAGAACAGCA |
| Cd36-RT-R | GGCTCAAAGATGGCTCC |
| Gpnmb-RT-F | AGAAATGGAGCTTTGTCTACGTC |
| Gpnmb-RT-R | CTTCGAGATGGGAATGTATGCC |
| Cd44-RT-F | GACCCAAGACTGTCACAGCA |
| Cd44-RT-R | AGTGGGCAATCCCTGTTCTT |
| GDF15-RT-F | TTCCTGCTGCTGTTGCT |
| GDF15-RT-R | TCCTCTCGGCTCTGGTT |
| TREM2-RT-F | ATGCCAGCGTGTGGTGA |
| TREM2-RT-R | GACGGTTCCAGCAAGGG |
| Lyz2-RT-F | TGTGAATGCCTGTGGGA |
| Lyz2-RT-R | AGACTCCGCAGTTCCGA |
| Fcgr1-RT-F | ACCTGAGTCACAGCGGCATCTA |
| Fcgr1-RT-R | TGACACGGATGCTCTCAGCACT |
| Csf3r-RT-F | CCTGGATGATAGAACCTAACGGG |
| Csf3r-RT-R | CTCTCCAGCGAAGGTGTAGACA |
| Fabp2-RT-F | TCCCTACAGTCTAGCAGACGGA |
| Fabp2-RT-R | CCAGAAACCTCTCGGACAGCAA |
| Irf7-RT-F | GAGTTTCGGGCTCGGAG |
| Irf7-RT-R | GGACACACCCTCACGCT |

| | |
|---|---|
| Fads2-RT-F | CAGAAGCACAACCTGCG |
| Fads2-RT-R | GGAAGGCATCCGTAGCA |
| Mmp12-RT-F | TTTGATGAGGCAGAAACGTG |
| Mmp12-RT-R | AGAGAGGCGAAATGTGCT |
| Mmp19-RT-F | TCCAGGCTCTCTATGGCA |
| Fas-RT-F | ATCTGGGCTGTCCTGCCT |
| Fas-RT-R | AGTTTCACGAACCCGCCT |
| Fabp1-RT-F | GCAATAGGTCTGCCCGAG |
| Fabp1-RT-R | TCCAGTTCGCACTCCTCC |
| Trem2-RT-F | CCTGAAGAAGCGGAATGGG |
| Trem2-RT-R | TCCAGCACCTCCACCAGTA |
| Lgals3-RT-F | AACACGAAGCAGGACAATAACTGG |
| Lgals3-RT-R | GCAGTAGGTGAGCATCGTTGAC |
| CD9-RT-F | TGCTGGGATTGTTCTTCGGG |
| CD9-RT-R | GCTTTGAGTGTTTCCCGCTG |
| Ccl6-RT-F | AAGAAGATCGTCGCTATAACCCT |
| Ccl6-RT-R | GCTTAGGCACCTCTGAACTCTC |
| Adgre1 (F4/80)-RT-F | ACCACAATACCTACATGCACC |
| Adgre1 (F4/80)-RT-R | AAGCAGGCGAGGAAAAGATAG |
